# Identifying perturbations that boost T-cell infiltration into tumours via counterfactual learning of their spatial proteomic profiles

**DOI:** 10.1101/2023.10.12.562107

**Authors:** Zitong Jerry Wang, Abdullah S. Farooq, Yu-Jen Chen, Aman Bhargava, Alexander M. Xu, Matt W. Thomson

## Abstract

Cancer progression can be slowed down or halted via the activation of either endogenous or engineered T cells and their infiltration of the tumour microenvironment. Here we describe a deep-learning model that uses large-scale spatial proteomic profiles of tumours to generate minimal tumour perturbations that boost T-cell infiltration. The model integrates a counterfactual optimization strategy for the generation of the perturbations with the prediction of T-cell infiltration as a self-supervised machine-learning problem. We applied the model to 368 samples of metastatic melanoma and colorectal cancer assayed using 40-plex imaging mass cytometry, and discovered cohort-dependent combinatorial perturbations (CXCL9, CXCL10, CCL22 and CCL18 for melanoma, and CXCR4, PD-1, PD-L1 and CYR61 for colorectal cancer) that support T-cell infiltration across patient cohorts, as confirmed via in vitro experiments. Leveraging counterfactual-based predictions of spatial omics data may aid the design of cancer therapeutics.

## Introduction

The immune composition of the tumor microenvironment (TME) plays a crucial role in determining patient prognosis and response to cancer immunotherapies [1–3]. Immunotherapies that alter the immune composition using transplanted or engineered immune cells (chimeric antigen receptor T cell therapy) or remove immunosuppressive signaling (checkpoint inhibitors) have shown exciting results in relapsed and refractory tumors in hematological cancers and some solid tumors. However, effective therapeutic strategies for most solid tumors remain limited [4–6]. The TME is a complex mixture of immune cells, including T cells, B cells, natural killer cells, and macrophages, as well as stromal cells and tumor cells [1]. The interactions between these cells can either promote or suppress tumor growth and progression, and ultimately impact patient outcomes. For example, high levels of tumor-infiltrating lymphocytes (TILs) in the TME are associated with improved prognosis and response to immunotherapy across multiple cancer types [7, 8]. Conversely, an immunosuppressive TME characterized by low levels of TILs is associated with poor prognosis and reduced response to immunotherapy [9]. Durable, long-term clinical response of T-cell-based immunotherapies are often constrained by a lack of T-cell infiltration into the tumor, as seen in classically “cold” tumors such as triple-negative breast cancer or pancreatic cancer, which have seen little benefit from immunotherapy [10–12]. The precise cellular and molecular factors that limit T-cell infiltration into tumors is an open question. Spatial omics technologies capture the spatial organization of cells and molecular signals in intact human tumors with unprecedented molecular detail, revealing the relationship between localization of different cell types and tens to thousands of molecular signals [13]. T-cell infiltration is modulated by a rich array of signals within the tumor microenvironment (TME) such as chemokines, adhesion molecules, tumor antigens, immune checkpoints, and their cognate receptors [14]. Recent advances in *in situ* molecular profiling techniques, including spatial transcriptomic [15, 16] and proteomic [17, 18] methods, simultaneously capture the spatial relationship of tens to thousands of molecular signals and T cell localization in intact human tumors with micron-scale resolution. Imaging mass cytometry (IMC) is one such technology that uses metal-labeled antibodies to enable simultaneous detection of up to 40 antigens and transcripts in intact tissue [17].

Recent work on computational methods as applied to multiplexed tumor images have primarily focused on predicting patient-level phenotypes such as survival, by identifying spatial motifs from tumor microenvironments [19–22]. These methods have generated valuable insights into how the complex composition of TMEs influences patient prognosis and treatment response, but they fall short of generating concrete, testable hypotheses for therapeutic interventions that may improve patient outcomes. Given the prognostic value of T-cell infiltration into tumors, we need computational tools that can predict immune cell localization from environmental signals and systematically generate specific, feasible tumor perturbations that are predicted to alter the TME to improve patient outcomes.

Counterfactual explanations (CFEs) can provide important insight in image analysis applications [23], but have not been applied to multiplexed imaging data. Traditionally, CFEs help clarify machine learning model decisions by exploring hypothetical scenarios, showing how the model’s interpretation would change if a feature in an image were altered slightly [24]. For instance, slight pixel intensity variations or minor edge alterations in a tumor’s appearance on an X-ray might lead a diagnostic model to classify the scan differently. Numerous CFE algorithms exist to elucidate a model’s decision boundaries and shed light on its sensitivity to specific image features [25]. In multiplexed tissue images where each pixel captures detailed molecular information, variations in pixel intensity directly correspond to specific molecular interventions. Thus, spatial omics data enables the extension of CFEs from understanding to predicting actionable interventions.

In this work, we introduce Morpheus, an integrated deep learning framework that first leverages large scale spatial omics profiles of patient tumors to formulate T-cell infiltration prediction as a self-supervised machine learning (ML) problem, and combines this prediction task with counterfactual optimization to propose tumor perturbations that are predicted to boost T-cell infiltration. Specifically, we train a convolutional neural network to predict T-cell infiltration using spatial maps of the TME provided by IMC. We then apply a gradient-based counterfactual generation strategy to the infiltration neural network to compute changes to the signaling molecule levels that increase predicted T-cell abundance. We apply Morpheus to melanoma [26] and colorectal cancer (CRC) with liver metastases [27] to discover tumor perturbations that are predicted to support T cell infiltration in tens to hundreds of patients. We provide further validation of ML-based T-cell infiltration prediction using an additional breast cancer data set [28]. For patients with melanoma, Morpheus predicts combinatorial perturbation to the CXCL9, CXCL10, CCL22 and CCL18 levels can convert immune-excluded tumors to immune-inflamed in a cohort of 69 patients. For CRC liver metastasis, Morpheus discovered two cohort-dependent therapeutic strategies consisting of blocking different subsets of CXCR4, PD-1, PD-L1 and CYR61 that are predicted to improve T-cell infiltration in a cohort of 30 patients. We experimentally validated Morpheus’ predictions by showing that perturbing these targets substantially enhanced T-cell migration *in vitro*. Our work provides a paradigm for counterfactual-based prediction and design of cancer therapeutics based on classification of immune system activity in spatial omics data.

## Results

### Counterfactual optimization for therapeutic prediction

The general logic of Morpheus (Methods, Figure 1A) is to first train, in a self-supervised manner, a classifier to predict the presence of CD8+ T cells from multiplexed tissue images (Figure 1B). Then we compute counterfactual instances of the data by performing gradient descent on the input image, allowing us to discover perturbations to the tumor image that increases the classifier’s predicted likelihood of CD8+ T cells being present (Figure 1C). The altered image represents a perturbation of the TME predicted to improve T-cell infiltration. We mask CD8+ T cells from all images to prevent the classifier from simply memorizing T-cell expression patterns, guiding it instead to learn environmental features indicative of T-cell presence.

**Fig. 1:**
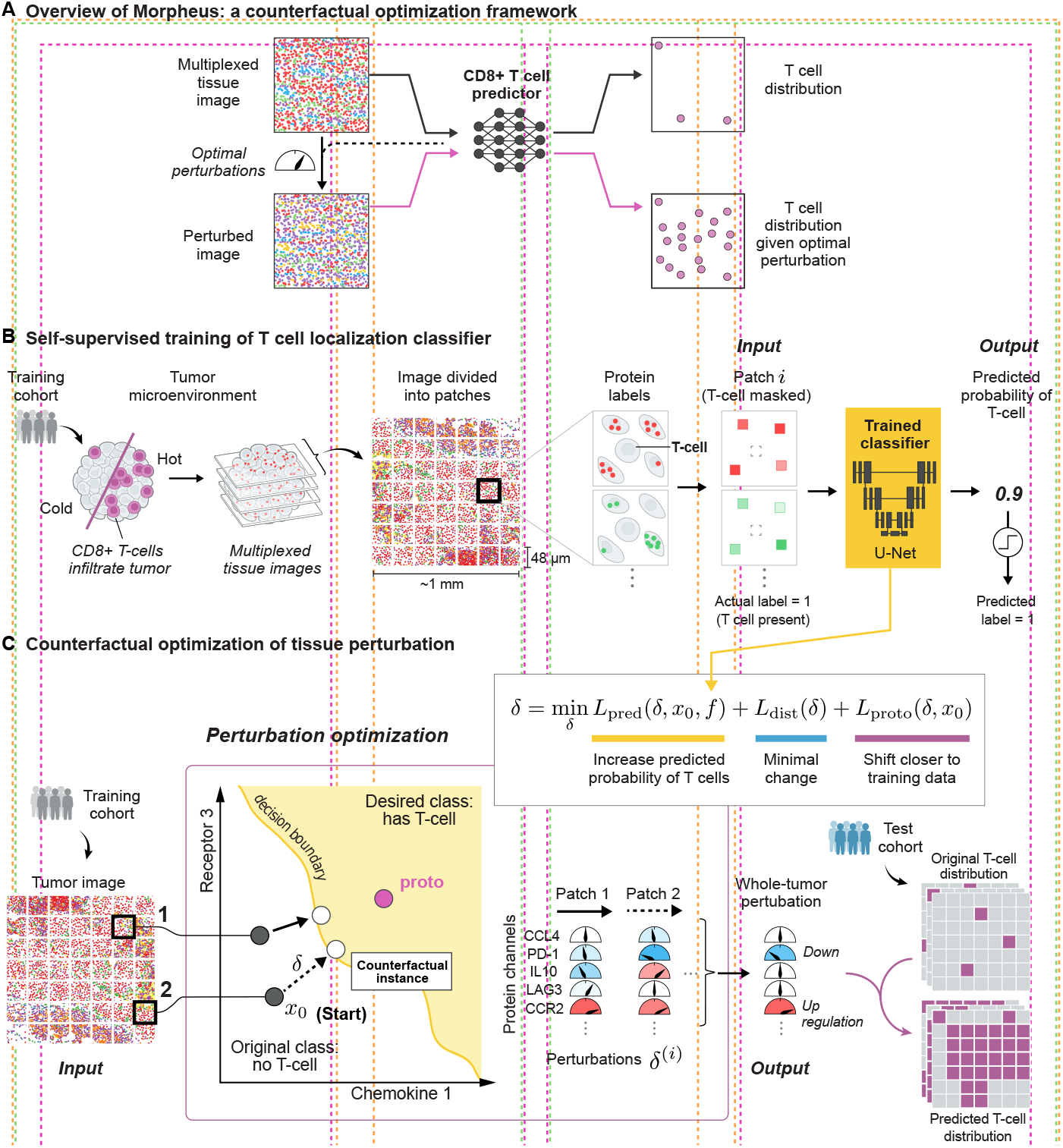
An integrated counterfactual optimization framework for discovering therapeutic strategies predicted to drive CD8+ T cell infiltration in human tumors. (A) Overview of the Morpheus framework, which consists of first (B) Training a neural network classifier to predict the presence of CD8+ T cells from multiplexed tissue images where cells in the IMC images are pixelated and CD8+ T cells are masked (Methods). (C) The trained classifier is then used to compute an optimal perturbation vector *δ*^(*i*)^ per patch by jointly minimizing three loss terms (*L*_pred_, *L*_dist_, *L*_proto_). The perturbation *δ*^(*i*)^ represents a strategy for altering the level of a small number of signaling molecules in patch 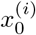 in a way that increases the probability of T cell presence as predicted by the classifier. The optimization also favors perturbations that shift the image patch to be more similar to its nearest T-cell patches in the training data, shown as proto. Each perturbation corresponds to adjusting the relative intensity of each imaging channel. Taking the median across all perturbations produces a whole-tumor perturbation strategy, which we assess by perturbing *in silico* tumor images from a test patient cohort and examining the predicted T cell distribution after perturbation.

We leverage IMC profiles of human tumors to train a model to predict the spatial distribution of CD8+ T cell in a self-supervised manner. We first divide IMC images into patches representing local tissue signaling environments, then we create a masked copy of each patch by removing all signals originating from CD8+ T cells (Figure 1B). We train a neural network model to classify whether T cells are present or absent using only the masked copy. Using our trained model, we apply counterfactual optimization to generate tumor perturbations predicted to enhance CD8+ T cell infiltration (Figure 1C). For each image patch *x*_0_ that does not contain CD8+ T cells, our optimization algorithm searches for a perturbation *δ* such that our classifier *f* predicts the perturbed patch *x*_*p*_ = *x*_0_ + *δ* as having T cells, hence *x*_*p*_ is referred to as a counterfactual instance. Furthermore, our algorithm favors simple and realistic strategies by minimizing the number of molecules perturbed while also ensuring the counterfactual instance is not far from image patches in our training data, so we can be more confident of the model’s prediction. We can obtain a perturbation *δ* with these desired properties by solving a constrained optimization problem (Methods).

Since drug treatments cannot act at the spatial resolution of individual micron-scale pixels, we constrain our search space to only perturbations that affect all cells in the image uniformly. Specifically, we only search for perturbations that change the level of any molecule by the same relative amount across all cells in an image.

Taken together, our algorithm obtains an altered image predicted to contain T cells from an original image which lacks T cells, by minimally perturbing the original image in the direction of the nearest training patch containing T cells until the classifier predicts the perturbed image to contain T cells. Since our strategy may find different perturbations for different tumor patches, we reduce the set of patch-wise perturbations {*δ*^(*i*)^}_*i*_ to a whole-tumor perturbation by taking the median across the entire set (Figure 1C).

### Convolutional neural networks predict T-cell distribution

We applied Morpheus to two publicly available IMC data sets of tumors from patients with metastatic melanoma [26] and colorectal cancer (CRC) with liver metastases [27] (Figure 2A). We validate the infiltration prediction on an additional breast cancer data set [28]. While this breast cancer data focuses on cell type markers over functional modulators of T-cell infiltration, making it unsuitable for therapeutic prediction, it serves to further validate our ML-based prediction of T-cell infiltration.

**Fig. 2:**
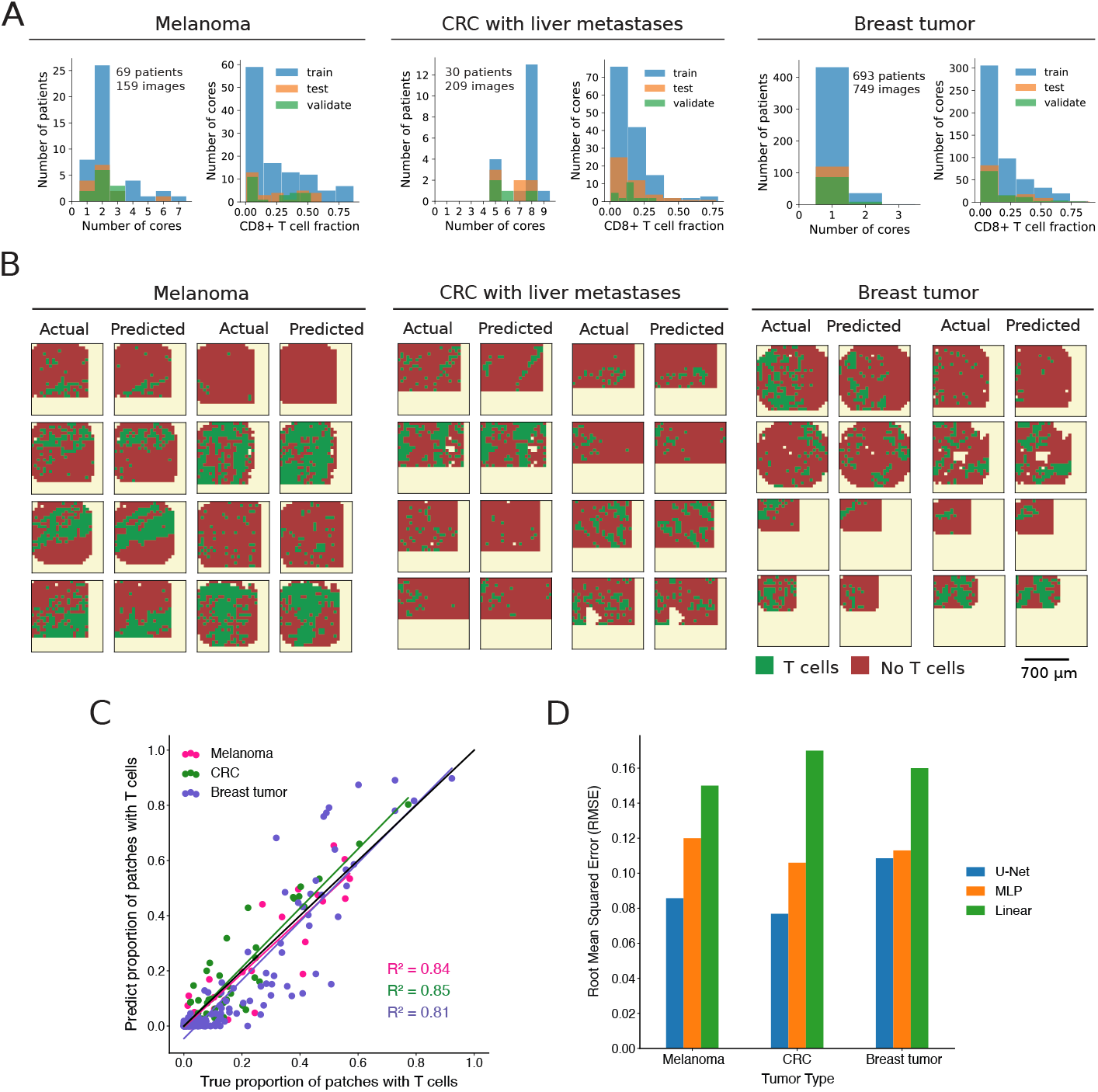
U-Net classifiers accurately predict T cell distribution in IMC images of melanoma, metastatic liver, and breast tumor. (A) Histograms showing the distribution of tumor cores per patient and CD8+ T cell fractions per core across all three data sets and data splits. (B) Predicted and actual T cell distribution of tissue sections from test cohorts in melanoma, liver tumor, and breast tumor data set. (C) Predicted and true proportion of patches with T cells within a tissue section, each dot corresponds to a tissue section, diagonal black line indicates perfect prediction. (D) The RMSE (Equation 3, Methods) across all (test) tissue sections for three different classes of models.

The melanoma data set [26] was obtained by IMC imaging of 159 tumor cores from 69 patients with stage III or IV metastatic melanoma. Each tissue was imaged across 39 molecular channels, consisting of markers for tumor, immune, and stromal cells, as well as 11 different chemokines (RNA) (Methods). The CRC data set [27] consists of 209 tissue sections taken from 30 patients imaged across 42 channels, including 60 sections from primary CRC tumors, 89 sections CRC metastases to the liver and 60 “healthy” liver sections obtained away from the metastases (Methods). The breast cancer data set [28] was obtained by IMC imaging of 749 breast tumor cores from 693 patients. The tissues were imaged across 37 channels, consisting of markers for tumor, lymphoid, myeloid and stromal cells (Methods).

For each of the three tumor data sets, we trained a separate U-Net classifier that effectively predicts CD8+ T cell infiltration level in unseen tumor sections (Methods). The two classifiers trained on melanoma and CRC data sets achieved the best performance (Supplemental Table 4) with an AUROC (area under the receiver operating characteristic curve) of 0.87 and 0.89 respectively, whereas the classifier trained on breast tumors achieved a AUROC of 0.83 (Supplemental Figure 2). A limited overlap in imaging channel across the three data sets makes it difficult to compare the TME across cancer types or to determine how difference in TME between the three cancer types affect Morpheus’ ability to predict T-cell presence. Figure 2B shows examples of actual and predicted T cell distributions in tumor sections, demonstrating that our classifiers accurately predict the general distribution of T cells. For each tissue section of a cancer type, the predictions were obtained by applying the corresponding U-Net classifier to each image patch independently. Comparing the true proportion of T-cell patches in a tissue section against our predicted proportion also shows strong agreement (Figure 2C). The true proportion of patches with T cells is calculated by dividing the number of patches within a tissue section that contain CD8+ T cells by the total number of patches within that section. We quantify the performance of our U-Nets on the entire test data set using the root mean square error (RMSE) (Equation 3, Methods), which represents the mean difference between our predicted proportion and the true proportion per tumor section (Figure 2D). Our classifiers performs well on liver tumor and melanoma, achieving a RMSE of only 7% and 8% respectively and a relatively poorer performance of 11% on breast tumor. Taken together, these results suggest that our classifier can accurately predict the T cell infiltration status of multiple tumor types.

In order to gain insight into the relative importance of non-linearity and spatial information in the performance of the U-Net on the T cell classification task, we compared the U-nets’ performance to a logistic regression model (LR) and a multi-layer perceptron (MLP). Both the LR and MLP model are given only mean channel intensities as input, so neither have explicit spatial information. Furthermore, the LR model is a linear model with a threshold whereas the MLP is a non-linear model. Figure 2D shows that across all three cancer data sets, the MLP classifier consistently outperforms the logistic regression model, reducing RMSE by 20 − 40% to suggest that there are considerable nonlinear interactions between different molecular features in terms of their effect on T cell localization. The importance of spatial features on the T cell prediction task, however, is less consistent across cancer types. Figure 2D shows that for predicting T cells in breast tumor, the U-Net model offers negligible boost in performance relative to the MLP model (*<* 2% RMSE reduction), whereas for liver tumor, the U-Net model achieved a RMSE 30% lower compared to the MLP model. This result suggests that the spatial organization of signals may have a stronger influence on CD8+ T cell localization in liver tumor compared to breast tumor.

### Applying Morpheus to metastatic melanoma samples

Applying our counterfactual optimization procedure using the U-Net classifier trained on melanoma IMC images, we discovered a combinatorial therapy predicted to be highly effective in improving T cell infiltration in patients with melanoma. Currently, there are substantial efforts to augment T-cell therapy using chemokines [29], which are a family of secreted proteins that are known for their ability to stimulate cell migration [30]. Since this data set is unique in its broad coverage of chemokine profiles, we applied Morpheus to systematically search for optimal chemokine therapy by restricting the optimization algorithm to only perturb chemokines. By optimizing over multiple chemokines, Morpheus opens the door to combinatorial chemokine therapeutics that has the potential to more effectively enhance T cell infiltration into tumors. Figure 3A shows that patients from the training cohort separate into two clusters based on hierarchical clustering of perturbations computed for each patient. Taking median across all patients in cluster 1, the optimized perturbation is to increase CXCL9 level by 215%, whereas in patient cluster 2, the optimized perturbation consists of increasing CXCL10 level by 88% while decreasing CCL18 and CCL22 levels by 100% and 100% respectively (Figure 3A). Both CXCL9 and CXCL10 are well-known for playing a role in the recruitment of CD8+ T cells to tumors. On the other hand, CCL22 is known to be a key chemokine for recruiting regulatory T cells [31] and CCL18 is known to induce an M2-macrophage phenotype [32], so their expression likely promotes an immunosuppressive microenvironment inhibitory to T cell infiltration and function.

**Fig. 3:**
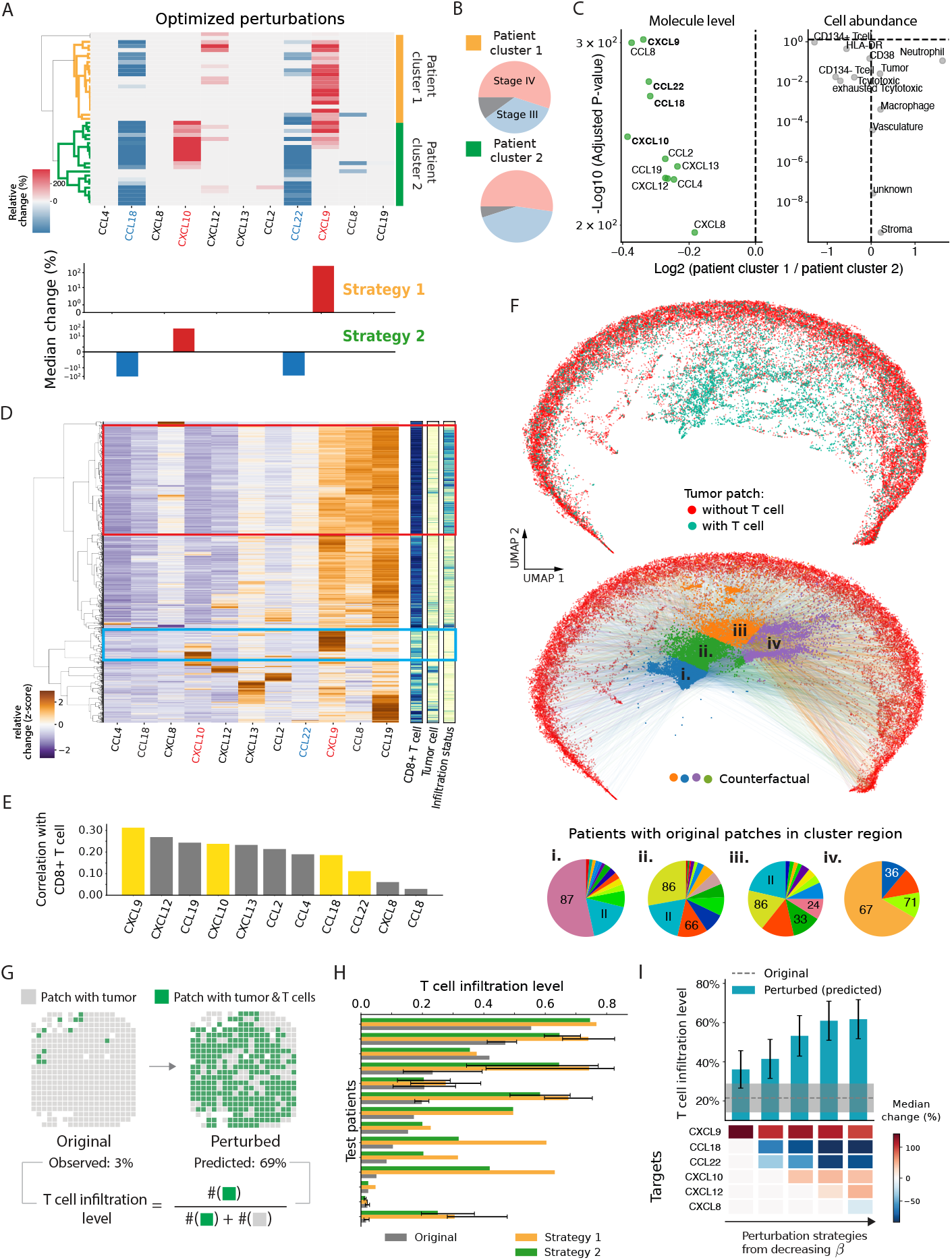
Combinatorial chemokine therapy predicted to drive T cell infiltration in patients with metastatic melanoma. (A) Whole-tumor perturbations optimized across IMC images of patients (row) from the training cohort, with bar graph showing the median relative change in intensity for each molecule. (B) Distribution of cancer stages among patients within two clusters, gray indicates unknown stage. (C) Volcano plot comparing chemokine level and cell type abundance from patient cluster 1 and 2, computed using mean values and Wilcoxon rank sum test with Šidák correction. Gray indicates non-statistical significance. Non-significant chemokines not shown: CXCL12 (FC = 0.96, *p* = 1) and CCL8 (FC = 0.93, *p* = 0.91) (D) Patch-wise chemokine profile (left); 1-D heatmap (right): infiltration status (light/dark = from infiltrated/deserted tumor), tumor cell (light/dark = present/absent), CD8+ T cells (light/dark = present/absent). (E) Patch-wise correlation between chemokine signals and the presence of CD8+ T cells. (F) (Top) UMAP projection of tumor patches (chemokine channels) show a clear separation of masked patches with and without T cells. (Bottom) colored arrows connect UMAP projection of patches without T cells and their corresponding counterfactual (perturbed) patch, where the colors correspond to k-nearest neighbor clusters (i-iv) of the counterfactual patches. Pie charts (iiv) shows the distribution of patients whose original tumor patches are found in the corresponding cluster regions in the UMAP. (G) Cell maps computed from a patient’s IMC image, showing the distribution of T cells before and after perturbation. (H) Original vs. perturbed (predicted) mean infiltration level across all patients (test cohort) with 95% confidence interval (only shown for patients with more than 2 samples). (I) Mean infiltration level across all patients (test cohort) for optimized perturbation strategies of varying sparsity, error bar represents 95% CI.

Figure 3B shows that the choice of which of these two strategies does not appear to be related to a patient’s cancer stage. We do find, however, nearly all chemokines have higher mean expression in the tumors of patients in cluster 2 compared to cluster 1, while there are no significant differences between the two groups in terms of the cell type compositions within tumors (Figure 3C). Since the levels of CCL22 and CCL18 is nearly 25% higher in patients from cluster 2 and both chemokines have been implicated in having an inhibitory effect on T-cell infiltration, it is reasonable that the optimization algorithm suggests inhibiting CCL18 and CCL22 only for patients in cluster 2. However, the switch from boosting CXCL9 to CXCL10 is not as straightforward. A possible explanation is that boosting CXCL10 is important when blocking CCL18 and CCL22 in order for the perturbed patches to stay close to the data manifold, leading to more realistic tissue environments. Interestingly, the single cell nature of the data set appears to be necessary for discovering this strategy as counterfactuals generated using pseudobulk data led to different strategies (Supplemental Note 3, Extended Data Figure 2A).

Morpheus selected perturbations that would make the chemokine composition of a TME more similar to T cell rich regions of immune-infiltrated tumors. Figure 3D shows that melanoma tissue patches can be clustered into distinct groups based on their chemokine concentration profile. One cluster (highlighted in blue) contains exactly the patches from immune-infiltrated tumors that contain both tumor and T cells, which likely represents a chemokine signature that is suitable for T cell infiltration. Alternately, a second cluster (highlighted in red) which contains patches from immune-desert tumors that have tumor cells but no T cells likely represents an unfavorable chemokine signature. In comparison to the cluster highlighted in red, Figure 3D shows the cluster highlighted in blue contains elevated levels of CXCL9, CXCL10 and reduced levels of CCL22 which partially agrees with the perturbation strategy (Figure 3A) discovered by Morpheus. Lastly, Figure 3E shows that our four selected chemokine targets cannot simply be predicted from correlation of chemokine levels with the presence of CD8+ T cells, as both CCL18 and CCL22 are weakly correlated (*<* 0.2) with CD8+ T cells even though the optimized perturbations requires inhibiting both chemokines, suggesting the presence of notable nonlinear effects not captured by correlations alone.

We can directly observe how Morpheus searches for efficient perturbations by viewing both the original patch and perturbed patches in a dimensionally-reduced space. Figure 3F (top) shows a UMAP (Uniform Manifold Approximation and Projection) projection where each point represents the chemokine profile of an IMC patch. T-cell patches (with their CD8+ T cells masked) are well-separated from patches without CD8+ T cells. The colored arrows in the bottom UMAP of Figure 3F illustrate the perturbation for each patch as computed by Morpheus, and demonstrate two key features of our algorithm. First, optimized perturbations push patches without T cells towards the region in UMAP space occupied by T-cell-infiltrated patches. Second, the arrows in Figure 3C are colored to show that optimized perturbations seem efficient in that patches are perturbed just far enough to land in the desired region of space. Specifically, red points that start out on the right edge end up closer to the right after perturbation (region iii and iv), while points that start on the left/bottom edge end up closer to the left/bottom (region i), respectively. We make this observation while noting that UMAP, though designed to preserve the topological structure of the data, is not a strictly distance-preserving transformation [33]. Furthermore, the pie charts (i-iv) are colored by the patient of origin to show the region of space where points are being perturbed to are not occupied by tissue samples from a single patient with highly infiltrated tumor. Rather, these regions consist of tissue samples from multiple patients, suggesting that our optimization procedure can synthesize information from different patients when searching for therapeutic strategies.

After applying the second perturbation strategy from Figure 3A *in silico* to IMC images of a tumor, Figure 3G shows that T cell infiltration level (defined as the proportion of tumor patches with T cells) is predicted to increase by 20 fold. In this data set, patients that respond favorably to immunotherapy tend to have significantly higher levels of T-cells within tumors before treatment (t-test, p = 0.006; Supplemental Figure 3). We applied both perturbation strategies on patients in our test cohort *in silico* and show that this predicted improvement holds across nearly all 14 patients from the test group, boosting T cell infiltration level from an average of 21% across samples to a predicted 50% post perturbation (Figure 3H).

The combinatorial nature of our optimized perturbation strategy is crucial to its predicted effectiveness. We systematically explored the importance of combinatorial perturbation by changing parameter *β* of equation (4) which adjusts the sparsity of the strategy, where a more sparse strategy means fewer molecules are perturbed. Figure 3I shows that perturbing multiple targets is predicted to be necessary for driving significant T cell infiltration across multiple patients, with the best perturbation strategy involving one target predicted to achieve 30% less T-cell level compared to the optimal strategy involving four targets. In conclusion, within the scope of the chemokine targets considered, combinatorial perturbation of the TME appears necessary for improving T cell infiltration in metastatic melanoma.

### Applying Morpheus to CRC with liver metastases samples

Applying Morpheus to IMC images from the CRC cohort, we discovered two patient-dependent therapies predicted to be highly effective in improving T cell infiltration (Figure 4A). Taking median over patients in the first cluster, the optimized strategy involves inhibiting PD-1, PD-L1, and CXCR4. While for the second group of patients, the optimized strategy involves inhibiting CYR61, PD-1, PD-L1, and CXCR4 (Figure 4A). Interestingly, all four of the perturbation targets correlated poorly with the presence of CD8+ T cells compared to the other proteins that were not selected as perturbation targets (Figure 4B), suggesting the presence of substantial spatial and nonlinear effects not captured by correlations alone.

**Fig. 4:**
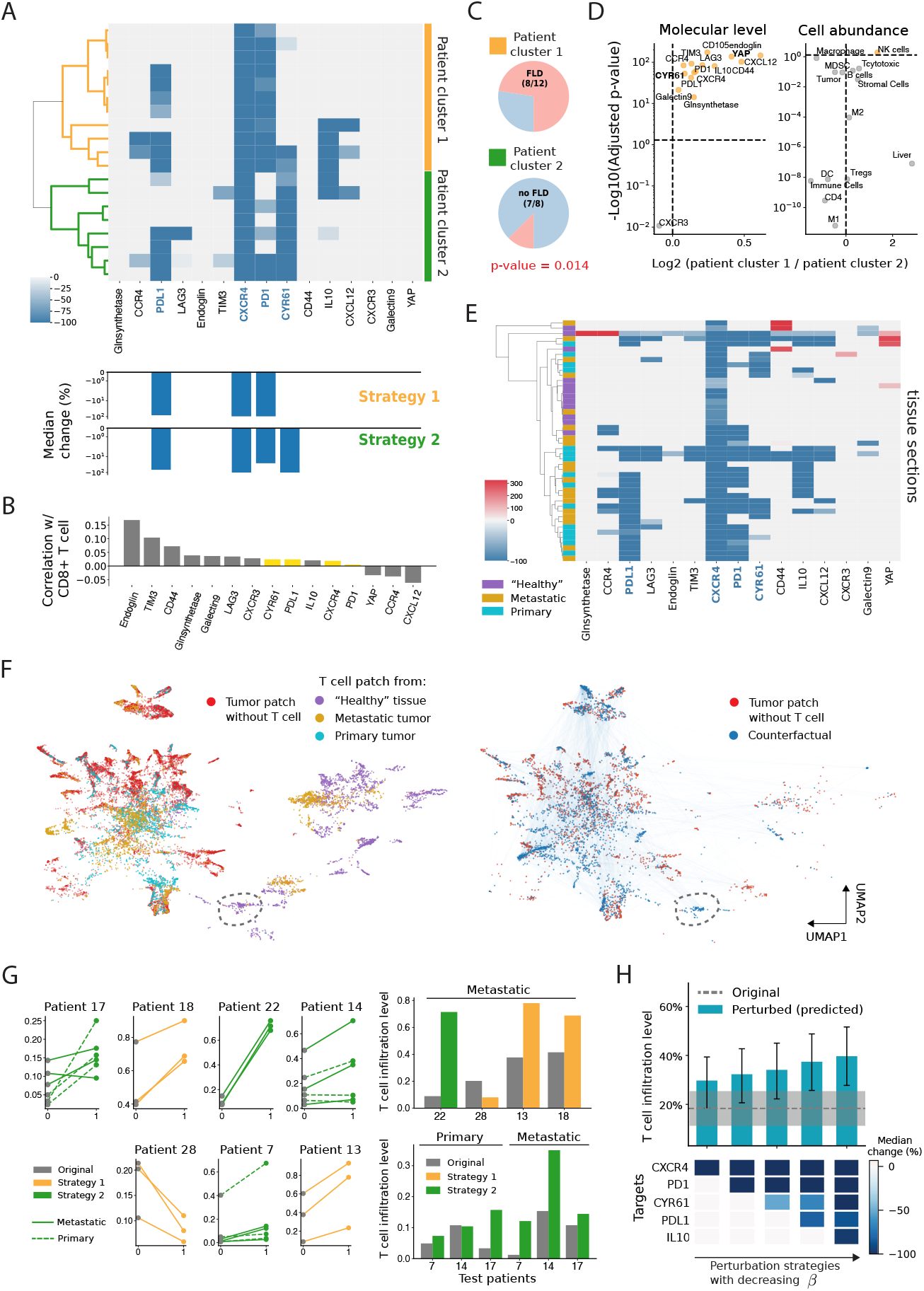
Blocking subsets of PD-L1, CXCR4, PD-1, and CYR61 predicted to drive T cell infiltration in CRC cohort. (A) Optimized tumor perturbations aggregated to the patient (row) level (train cohort). Bar graph shows the median relative change in intensity for each molecule across all patients within their cluster. (B) Patch-wise correlation between the levels of different molecules and the presence of CD8+ T cells. (C) Pie charts show proportion of patients in each cluster that have fatty liver disease (FLD), p-value from hypergeometric test. (D) Volcano plot comparing molecule levels and cell type abundance between the two patient cluster using tumor tissues, computed using mean values and Wilcoxon rank sum test with Šidák correction. (E) Optimized perturbations aggregated to the level of tissue samples (row). (F) UMAP projection of IMC patches, left UMAP shows T cell patches colored by the tissue samples they are taken from. right UMAP shows counterfactual (perturbed) instances optimized for tumor patches without T cells (red). (G) Line plots shows T-cell infiltration level for each tissue section from the test cohort, before and after perturbation. Bar plots show predicted mean T-cell infiltration level for each test patient. (H) Mean infiltration level across all test patients using perturbation strategies of varying sparsity, obtained by varying *β* in equation (4), error bar represents 95% CI.

All perturbation targets identified by our optimization procedure have been found to play crucial roles in suppressing T cell function in the TME, and treating patients with inhibitors against subsets of the selected targets have been shown to improve T cell infiltration in human CRC liver metastases. Tumor-associated lymphatic vessels promote T cell exit from tumor via the CXCL12/CXCR4 axis [34], and the PD-1/PD-L1 pathway inhibits CD8+ T cell activity and infiltration in tumors. In addition, CYR61 is a chemoattractant and was recently shown to drive M2 TAM (tumor-associated macrophage) infiltration in patients with CRC liver metastases [27]. Inhibition of both PD-1 and CXCR4, which were consistently selected by Morpheus as targets, have already been shown to increase CD8+ T cell infiltration in preclinical mouse models of colon cancer [35, 36]. The single cell nature of the CRC data set appears to be necessary for discovering this strategy as counterfactuals generated using pseudobulk data led to different strategies (Supplemental Note 3, Extended Data Figure 2B).

The emergence of the two distinct perturbation strategies may be explained by variation in liver fat build-up among patients. Patient cluster 1 is made up of significantly more patients with FLD or fatty liver disease (67%) compared to patient cluster 2 (12%) (Figure 4C). Furthermore, Figure 4D shows that both YAP and CYR61 levels are significantly higher in tumors from patient cluster 1, by 50% and 3.5% respectively. Indeed, CYR61 is known to be associated with non-alcoholic fatty liver disease [27] and YAP is a transcription coregulator that induces CYR61 expression [37]. However despite patients in cluster 1 having higher levels of CYR61, it is only for patients in cluster 2 where the optimal strategy involves blocking CYR61. We postulate that this seemingly paradoxical finding may arise because removing CYR61 from patients in cluster 1 represents a more pronounced perturbation, given their inherently higher concentration. A perturbation of this magnitude would likely shift the tumor profile substantially away from the data manifold, where the classifier’s prediction about the perturbation’s effect becomes less reliable, hence such a perturbation would be heavily penalized during optimization due to the *L*_proto_ term.

Using only raw image patches, Morpheus discovers tissue-dependent perturbation strategies (Figure 4E). As depicted in Figure 4E, by aggregating perturbations at the individual tissue level, we observe that the optimized perturbation for “healthy” liver sections is straightforward, necessitating only the inhibition of CXCR4. Recall “healthy” sections are samples obtained away from sites of metastasis. In contrast, promoting T cell infiltration into primary colon tumors is anticipated to involve targeting a minimum of three signals. Morpheus finds that liver metastases appears to fall between these two tissue types. Furthermore, direct comparison between perturbations optimized for metastatic tumor and primary tumor samples does not reveal a notable difference in strategy (Supplemental Figure 1). We can partly understand the discrepancy between tissues by plotting a UMAP projection of all T cell patches from the three tissue types (Figure 4F, left). The clear separation between T cell patches from “healthy” tissue and those from primary tumors underscores that the signaling compositions driving T cell infiltration likely differ substantially between the two tissue types, prompting Morpheus to identify markedly different perturbation strategies. Furthermore, some patches from metastatic tumors co-localize with “healthy” tissue patches in UMAP space, while other patches co-localizes with primary tumor patches. This observation again aligns with our previous result, where optimized perturbation strategies for metastases samples share similarities with strategies for either “healthy” tissue or primary tumor (Figure 4E).

Despite the CRC data set comprising a mixture of healthy, tumor, and hybrid metastatic samples, Morpheus targets the most pertinent tissue type when optimizing perturbations. During both the model training and counterfactual optimization phases, we did not make specific efforts to segregate the three tissue types. Furthermore, we did not provide tissue type labels or any metadata. Despite these nuances, Figure 4F shows that the counterfactual instances for tumor patches (dark blue) from primary and metastases samples are mostly perturbed to be near T cell patches from primary (cyan) and metastatic tumor (gold), instead of being perturbed to be similar to T cell patches from “healthy” tumors (purple). This result is partly a consequence of our prototypical constraint which encourages patches to be perturbed towards the closest T-cell patch. For a patch from a metastatic tumor without T cells, the closest (most similar) T cell patch is likely also from a metastatic tumor than from a “healthy” tissue. However, there are occasional exceptions where T cell patches from “healthy” tissues can influence the optimization of tumor tissues, as outlined by the dashed ellipse in Figure 4F, especially if they share similar features as tumor regions.

The two therapeutic strategies we discovered generalize to patients in our test cohort (Figure 4G,H). Given that we have two therapeutic strategies, one enriched for patients with FLD and another for patients without FLD, we apply different perturbation strategies *in silico* across all test patients depending on their FLD status. Aggregated to the patient level, Figure 4G shows that CD8+ T cell infiltration level is predicted to increase for nearly all patients, significantly boosting mean infiltration level from 17% to a predicted 35% post perturbation (Figure 4H). However, when comparing individual tissue samples, Figure 4G reveals substantial variation in the predicted response to perturbation among samples from the same patient and tissue types. In patient 7, two metastatic tumor sample is predicted to see a nearly seven-fold increase in T cell infiltration after perturbation, yet almost no change is expected for patient 7’s other three primary and one metastatic samples. Similar patterns are observed in patients 14 and 17. This marked variability in response among a substantial portion of test patients underscores the challenges posed by intra-tumor and inter-patient heterogeneity in devising therapies for CRC with liver metastases. This result further implies that, for studying CRC with liver metastases, collecting numerous tumor sections per patient could be as crucial as establishing a large patient cohort.

Lastly, combinatorial perturbation is again predicted to be necessary to drive significant T-cell infiltration in large patient cohorts. By increasing *β* in equation (4), we generated strategies with between one and five total targets, where perturbing at least four targets is predicted to be necessary to produce a statistically significant boost to T-cell infiltration (Figure 4H).

### Experimental validation of predicted perturbation strategies

Morpheus-derived strategies boost T cell level in *in vitro* migration assays using human melanoma and colorectal cancer cells. We tested Morpheus’ predictions using a transwell migration assay (Methods), which consists of two chambers separated by a permeable membrane allowing for selective passage of molecules and cells (Figure 5A). We place human peripheral blood mononuclear cells, PBMCs, initially in the top chamber and human cancer cells in the bottom chamber, where we apply different perturbations proposed by Morpheus and count the number of CD8+ T cells that infiltrate the bottom chamber after four hours using flow cytometry (Methods). This transwell/co-culture system is a common method for assessing the effect of different perturbations in altering the migratory capacity of T cells towards cancer cells. The PBMC cell population contains a mixture of immune cell types including CD8+ T-cells. For CRC, perturbations are applied to both chambers as some target molecules are expressed by immune cells (e.g. PD-L1, CXCR4). Similar transwell/co-culture systems are commonly used for assessing the effect of different perturbations in altering the migratory capacity of T cells towards cancer cells [38–40].

**Fig. 5:**
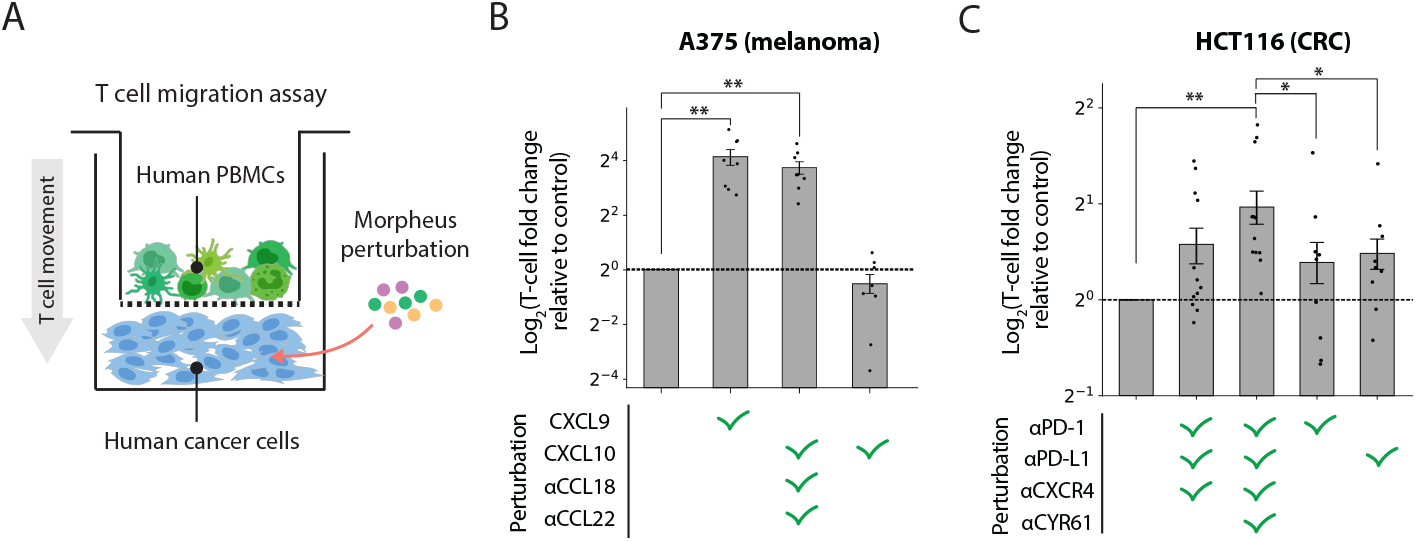
*in vitro* experimental validation of Morpheus predictions. (A) T-cell transwell migration assay for assessing the effect of Morpheus-derived perturbation strategies on CD8+ T-cell infiltration into an *in vitro* tumor compartment. Human PBMC cells are placed into the top chamber, and a human tumor cell line (A375 for melanoma, HCT116 for CRC) is placed into the bottom chamber. We measure CD8+ T-cell infiltration into the bottom chamber after four hours in the presence or absence of signaling perturbations predicted by Morpheus. Signaling perturbations include both signaling protein addition and blocking antibodies which are indicated by *α*/anti (B) Log fold change in CD8+ T-cell abundance within the lower chamber containing A375 melanoma cells relative to CD8+ T-cell abundance in unperturbed controls. CXCL9 and {CXCL10, *α*CCL18,*α*CCL22} are Morpheus predicted infiltration strategies while the CXCL10 addition alone strategy is shown for comparison to CXCL9 alone. (C) Log fold change in CD8+ T-cell abundance within the lower chamber containing HCT116 CRC cells relative to CD8+ T-cell abundance in unperturbed control. {*α*PD-1, *α*PD-L1, *α*CXCR4} and {*α*PD-1, *α*PD-L1, *α*CXCR4, *α*CYR61} are Morpheus predicted strategies. The *α*PD-1 and *α*PD-L1 strategy, blocking PD-1 or PD-L1, are clinical immunotherapy strategies shown for comparison. Each perturbation trial was normalized to its paired control trial. (*) indicates P-value *<* 5 × 10^−2^ and (**) indicates P-value *<* 1× 10^−2^. Two-sided paired t-tests used to assess significance, see Supplemental Table 6 and Supplemental Table 7 for raw data. The error bars represent the mean ±s.e.m. of independent biological replicates and dots indicate individual *n* = 8 and *n* = 9 replicate values for (B) and (C), respectively (*n* = 12 for bar 2-3 in (C)).

We used the human A375 melanoma cell line to test both sets of melanoma perturbations (Figure 3A). Directly adding either CXCL9 proteins alone or a triple strategy consisting of CXCL10, anti-CCL22 antibody, and anti-CCL18 antibody increased T-cell level in the tumor chamber by 17 and 14 fold, respectively, (paired t-test, p = 1×10^−3^ and 1×10^−4^, respectively; Figure 5B). Although CXCL9 and CXCL10 are often considered to have similar functions as they both bind to the receptor CXCR3 and act as chemoattractants for CD8+ T cells, Morpheus did not predict the addition of CXCL10 alone as an effective perturbation, rather Morpheus always predicted CXCL10 perturbation as one part of a combinatorial perturbation (Figure 3A). We found that, in fact, the addition of CXCL10 alone did not lead to any significant increase in T-cell level compared to the perturbed control (paired t-test, p = 0.09; Figure 5B).

We used the human HCT116 colorectal cell line to test both sets of CRC perturbations (Figure 4A), by adding either blocking antibodies against PD-1, PD-L1, and CXCR4 or an additional blocking antibody against CYR61. In close agreement with model predictions (Figure 4H), Morpheus’ four-target combinatorial perturbation increased T-cell abundance by 2-fold compared to unperturbed control (paired t-test, p = 2×10^−3^). The Morpheus four-target strategy also significantly outperforms anti-PD1 and anti-PD-L1 treatment alone in our *in vitro* assays where we observed 1.3-fold and 1.4-fold increase in T-cell abundance, for anti-PD1 and anti-PD-L1 respectively. The T-cell abundance change induced by anti-PD1 and anti-PD-L1, which represent standard immunotherapies, was significantly lower than that induced by Morpheus across replicates as quantified by a paired t-test with p = 0.02 for Morpheus four-target vs PD-1 and p = 0.04 for Morpheus four-target vs PD-L1 (Figure 5C). Unlike the four-target combination, we did not observe significant increase in T-cell infiltration with the three target strategy predicted by Morpheus (inhibition PD-1, PDL1, CXCR4) (1.5-fold increase, paired t-test, p = 0.09). We hypothesize that this relatively modest improvement from this three-target strategy is due to the absence of lymphatic endothelial cells in our *in vitro* assay. It was recently shown that tumor-associated lymphatic vessels control T cell exit from tumor through the interaction between CXCL12 and CXCR4, and inhibiting CXCR4 boosts the quantity of intratumoral T cells specifically in tumors with lymphatic vessel-derived CXCL12 [34].

Altogether, we show that experimentally perturbing molecular targets according to Morpheus’ predicted strategy consistently improves the ability of T cells to migrate towards cancer cells *in vitro*. For cancers for which PD-1/PD-L1 therapy is ineffective, Morpheus suggests new alternatives with promising *in vitro* results

## Discussion

Our integrated deep learning framework, Morpheus, combines deep learning with counterfactual optimization to directly predict therapeutic strategies from spatial omics data. One of the major strengths of Morpheus is that it scales efficiently to deal with large diverse sets of patients samples including metachronous tissue from the same patients but different sites, which will be crucial as more spatial transcriptomics and proteomics data sets are quickly becoming available [41].

Morpheus identifies fundamentally different strategies to increase T-cell abundance, beyond just enhancing the rate of T-cell entry into the tumor. In the literature, the term “infiltration” is often used as a catch-all term to refer to T-cell abundance. For clarity, while we align with this terminology, our focus is on strategies that boost overall T-cell abundance. For example, Morpheus’ strategy of inhibiting CXCR4 prevents T-cells from exiting the tumor via the vasculature, thereby increasing T-cell abundance by reducing outflow. This illustrates Morpheus’ ability to reveal diverse mechanisms for enhancing T-cell presence in tumors.

While the molecular targets identified by Morpheus require experimental validation to confirm their causal role in T-cell infiltration, both the biological context and Morpheus’ counterfactual optimization objective strengthen their potential value. Biologically, many effects of T-cells on their microenvironment feed back to influence further T-cell infiltration [42–44], creating cyclical relationships where effects may become causes. For example, T-cells promote tertiary lymphoid structures (TLS) supporting further infiltration [42], release cytokines like interferon-gamma (IFN-*γ*) to both inhibit and facilitate T-cell infiltration by inducing PD-L1 expression and upregulating chemokines [43], and enhance vascular permeability to facilitate additional T-cell infiltration [44]. Morpheus’ counterfactual optimization objective helps to address the association vs. causation challenge by focusing on minimal interventions with significant effects. This approach attempts to prioritize direct causal factors, which typically have stronger associations with outcomes than non-causal correlates. While these factors don’t guarantee causality, they may increase the likelihood that the identified targets meaningfully influence T-cell infiltration.

Nonetheless, further experimental validation remains necessary to confirm any causal relationships.

For future work, we would like to apply Morpheus to spatial transcriptomics data sets with hundreds to thousands of molecular channels. Although spatial transcriptomics can profile substantially more molecules compared to spatial proteomic techniques [15, 16], the number of spatial transcriptomic profiles of human tumors is currently limited due to the cost, with most public data sets containing single tissue sections from 1-5 patients which is far too small to apply Morpheus. However, spatial transcriptomics is likely to be more standardized compared to proteomics, which use customized panels. As commercial platforms for spatial transcriptomics start to come online [45], we will likely be seeing large scale spatial transcriptomics data sets in the near future, with ∼70-90% of the same probes shared between experiments.

A technical extension of Morpheus involves incorporating prior knowledge of gene-gene interactions to model the causal relations between genes. Molecular features in tissue profiles can exhibit strong dependencies, therefore, changing the level of one molecule can affect the expression of others. For example, increased levels of IFN-*γ* in the tumor microenvironment, can upregulate the expression of PD-L1 on tumor cells [46]. In order to be more realistic and actionable, a counterfactual should maintain these known causal relations. We can apply a regularizer to penalize counterfactuals that are less feasible according to established gene interactions from knowledge graphs, such as Gene Ontology [47].

Other extensions of Morpheus includes predicting cell-type specific perturbations, which can be done by directly restricting the perturbation to only alter signals within specific cell types. Additionally, although we applied Morpheus to the specific problem of driving T cells to infiltrate solid tumors, we can generalize our framework to predict candidate therapeutics that alter the localization of other cell types. For example, Morpheus can train a classifier model to predict localization of TAMs and compute perturbations predicted to reduce their abundance in the TME.

In this work, we focused on identifying generalized therapies by pooling predictions across multiple patient samples, but we can also apply Morpheus to find personalized therapy for treating individual patients. The variation in the optimized perturbations we observe among patients in both melanoma and liver data sets suggest personalize treatments could be substantially more effective compared to generalized therapies (Figure 3A, Figure 4A). Furthermore, Figure 4G shows that a therapeutic strategy could have highly variable effect even across different tissue samples from the same patient. This variability suggests that to generate therapy for an individual patient, it may be necessary to acquire substantial quantities of biopsy data. We can then apply our optimization procedure to a random subset of the samples, and then test the resulting perturbation strategy on the remaining samples to see how well the strategy is predicted to perform across an entire tumor or other primary/secondary tumors.

Incorporating Morpheus in a closed loop with experimental data collection is another promising direction for future work. Data can be collected from patients or animal models with perturbed/engineered signaling context, and this data can be easily fed back into the classifier model to refine the model’s prediction. The perturbation could be based on what the model predicts to be effective interventions, as is the case with Morpheus. We can also study tissue samples on which the model tends to make the most mistake and train the model specifically using samples from similar sources, such as similar patient strata or disease state.

## Methods

### Description of IMC data sets

All data sets used in this paper are publicly available. Metastatic melanoma data set from Hoch et al. [26] contains 159 images or cores taken from 69 patients, collected from sites including skin and lymph-node. CRC liver metastases data set from Wang et al. [27] contains 209 images or cores taken from 30 patients. Breast tumor data set from Danenberg et al. [28] contains 693 images or cores taken from 693 patients. The RNA and protein panels used for each of the three data sets are listed in Supplemental Table 8.

### Data split

For all three IMC data sets, we followed the same data splitting scheme to divide patients into three different groups (training, validation, testing) while ensuring similar class balance across the groups, which in our case means that the proportion of image patches with and without T cells are roughly equal across the three groups for each data set. Patients are shuffled between the three groups until three criteria are met: 1) the number of patients across the three groups follow a 65/15/20 ratio, 2) the difference in class proportion between any two of the three groups is less than 2%, and 3) the training set contains at least 65% of total patches. The actual data splits used in the paper are described in Supplemental Table 9.

### Overview of the Morpheus framework

The Morpheus framework consists of two main steps, the first being selfsupervised training of a classifier to predict the presence of CD8+ T cells from multiplexed tissue images (Figure 1B). Then we compute counterfactual instances of the data by performing gradient descent on the input image, allowing us to discover perturbations to the tumor image that increases the classifier’s predicted likelihood of CD8+ T cells being present (Figure 1C). The perturbed image corresponds directly to a perturbation of the TME predicted to improve T-cell infiltration. We mask CD8+ T cells from all images to prevent the classifier from simply memorizing T-cell expression patterns, guiding it instead to learn environmental features indicative of T-cell presence. We will describe both model training and counterfactual optimization in detail in the following sections.

### Training a classifier to predict T cell localization

We leverage IMC profiles of human tumors to train a classifier to predict the spatial distribution of CD8+ T cell in a self-supervised manner.

### Cell segmentation and phenotyping

Raw IMC images were processed to obtain single-cell masks using the ImcSegmentationPipeline [48]. The segmentation process began by converting raw data to ome-tiff format, followed by pixel classification in Ilastik [49], which segmented images into nuclear, cytoplasmic, and background regions. Probability maps generated from Ilastik were further processed in CellProfiler [50] to create single-cell masks. To correct for channel spillover, a non-negative least squares method was applied using CATALYST [51]. For cell phenotyping, we adopted an automated approach complemented by manual curation. Singlecell segmentation masks were overlaid with single-channel tiff images to extract mean marker expression values for each cell, which were then arcsinh transformed using a cofactor of 1 and censored at the 99th percentile. Cell clusters were determined by applying PhenoGraph [52] on the single-cell expression vectors using default hyperparameters. Channel gating was employed to refine the identification of specific cell populations. Two levels of PhenoGraph clustering were performed: the first level identified major cell types (immune, stromal and tumor) and the second level further classified immune cells into subtypes. For the purpose of Morpheus, only identification of CD8+ T cells and tumor cells was required. Manual curation of cell clusters was guided by specific proteins (Supplemental Table 10) to ensure accurate classification. See Supplemental Table 3 for cell type distribution.

### Cell pixelation of IMC images

The purpose of model training is for the model to learn molecular features of a tissue environment that supports the presence of CD8+ T cell, so it is important for us to remove features of the image that are predictive of CD8+ T cell presence but are not part of the cell’s environment, such as the expression of the T cell itself. A simple masking strategy of zeroing out all pixels belonging to CD8+ T cells will introduce contiguous regions of zeros to image patches with T cells, which is an artificial feature that is nonetheless highly predictive of T-cell presence and hence will likely be the main feature learned by a model during training. To circumvent this issue, we first apply a cell “pixelation” step to each IMC image, where we reduce each cell to a single pixel positioned at the cell’s centroid. The value of this pixel is the sum of all pixels originally associated with the cell, representing the total signal from each channel within the cell. In this way, we can simply mask this “pixelated” version of the image by zeroing all pixels representing CD8+ T cells. Our strategy is effective at masking T cells without introducing an artificial signal whereby simply removing cells at random will increase the chance that T cells are predicted to be present (Supplemental Note 1 Assessing effects of data pre-processing steps, Supplemental Table 1).

### Patching and T cell masking

From the set of “pixelated” IMC images, we obtain a set of image patches {*I*^(*i*)^} by first dividing each image into local patches of tissue and then downsample each patch using a max-pooling operation (3 ×3 kernel, stride = 3, no padding) to reduce the dimensionality of the input with minor information loss (Supplemental Table 2). Thus *I*^(*i*)^ ∈ℝ^*l×w×c*^ is an array with *l* and *w* denoting the pixel length and width of the image and *c* denoting the number of molecular channels in the images (Figure 1B). Each image patch shows the level of *c* proteins across all cells within a small region of tissue. We set *l* = *w* = 16, corresponding to a 48 µm ×48 µm region (previously each pixel = 1 µm, now each pixel = 3 µm due to downsampling). We applied spectral analysis to study the effect of using different patch size to predict T cell infiltration and found that our selected patch size remains highly informative of T cell presence (Supplemental Note 2, Extended Data Figure 1).

From a patch *I*^(*i*)^, we can obtain a binary label *s*^(*i*)^ indicating the presence and absence of CD8+ T cells in the patch and a masked copy *x*^(*i*)^ with all signals originating from CD8+ T cells removed (set to zero). The task for the model *f* is to classify whether T cells are present (*s*^(*i*)^ = 1) or absent (*s*^(*i*)^ = 0) in image *I*^(*i*)^ using only its masked copy *x*^(*i*)^. Specifically, *f* (*x*^(*i*)^) ∈[0, 1] is the predicted probability of T cells, and then we apply a classification threshold *p* to convert this probability to a predicted label *ŝ*^(*i*)^ ∈ {0, 1}. Since we obtain the image label *s*^(*i*)^ from the image *I*^(*i*)^ itself by unsupervised clustering of individual cell expression vectors, our overall task is inherently self-supervised.

### Classifier training objective

Given a set of masked image patches {*x*^(*i*)^} with corresponding CD8+ T cell label {*s*^(*i*)^}, we train a model *f* to minimize the following T cell prediction loss, also known as the binary cross entropy (BCE) loss,

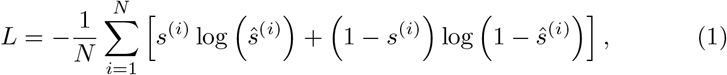

where

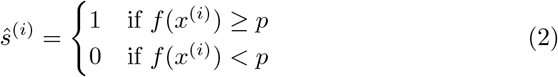

and *p* is the classification threshold. We select *p* by minimizing the following root mean squared error (RMSE) on a separate set of tissue sections Ω,

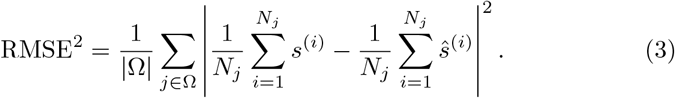

The RMSE is a measure of the differences between the observed and predicted proportions of T cell patches in a tissue section averaged across a set of tissues Ω, which we take to be the validation set.

### U-Net architecture

To obtain a model that can accurately predict T cell localization from environmental cues, we trained a fully convolutional neural network with the U-Net architecture to minimize (Equation 1). The U-Net architecture consists of a contracting path and an expansive path, which gives it a U-shaped structure [53]. The contracting path consists of four repeated blocks, each containing a convolutional layer followed by a Rectified Linear Unit (ReLU) activation and a max pooling layer. The expansive path mirrors the contracting path, where each block contains a transposed convolutional layer. Skip connections concatenates the up-sampled features with the corresponding feature maps from the contracting path to include local information. The output of the expansive path is then fed to a fully-connected layer with softmax activation to produce a predicted probability.

### U-Net training

We train our U-net classifiers on patches obtained from patients from the training cohort, using stochastic gradient descent with momentum and a learning rate of 10^−2^ on mini-batches of size 128. Image augmentation was used to prevent over-fitting, including random horizontal/vertical flips and rotations, in addition to standard channel-wise normalization. All models presented in this paper were trained with early stopping based on the validation Matthews Correlation Coefficient (MCC), computed using patches from the validation cohort, for a max of 30 epochs. All model performances are reported on patches from the test cohort. All models were trained on an NVIDIA GeForce RTX 3090 Ti GPU using PyTorch v2.0.0 [54] and PyTorch Lightning v2.2.2 [55]. Implementation code can be found in our Github repository along with a tutorial Jupyter notebook illustrating the entire workflow using an example data set.

We evaluated the performance of various classifiers, including both traditional convolutional neural networks (CNNs) and vision transformers. In all cases, we observed similar performance (Supplemental Table 5). We settled on a U-Net architecture because of ease of extension of the model to multichannel data sets.

### Generating counterfactuals using T cell prediction model

Our trained model allows us to formulate counterfactual optimization as a constrained optimization problem to generate tumor perturbations predicted to enhance CD8+ T cell infiltration (Figure 1C).

### Mathematical formulation of optimization problem

Given an image patch 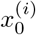 that does not contain CD8+ T cells, our optimization algorithm searches for a perturbation *δ*^(*i*)^ such that our classifier *f* predicts the perturbed patch 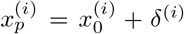 as having T cells, hence 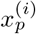 is referred to as a counterfactual instance. Ideally, we want each perturbation to involve perturbing as few molecules as possible, and realistic in that the counterfactual instance is not far from image patches in our training data so we can be more confident of the model’s prediction. We can obtain a perturbation *δ*^(*i*)^ with these desired properties by solving the following optimization problem adopted from [56],

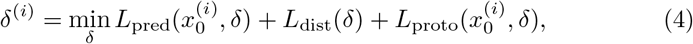

such that

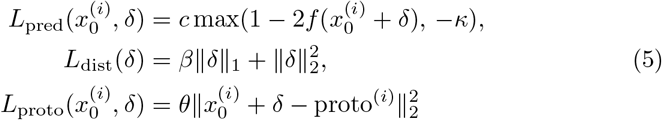

where *δ*^(*i*)^ is a 3D tensor that describes perturbation made to each pixel of the patch.

The three loss terms in equation (4) each correspond to a desirable property of the perturbation we aim to discover. The term *L*_pred_ encourages validity, in that the perturbation increases the classifier’s predicted probability of T cells, so the network is more likely to predict the perturbed tissue patch as having T cells when it previously did not contain T cells. Next, the term *L*_dist_ encourages sparsity using elastic net regularization, favoring perturbations that do not require making many changes to the TME. Lastly, the term proto^(*i*)^ in the expression for *L*_proto_ refers to the nearest neighbour of 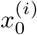 among all patches in the training set that are classified as having T cells. Thus the term *L*_proto_ explicitly guides the perturbed image 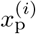 to lie close to the data manifold defined by our training set, making perturbed patches appear similar to what has been observed in TMEs infiltrated by T cells.

Since drug treatments cannot act at the spatial resolution of individual micron-scale pixels, we constrain our search space to only perturbations that affect all cells in the image uniformly. Specifically, we only search for perturbations that change the level of any molecule by the same relative amount across all cells in an image. We incorporate this constraint by defining *δ*^(*i*)^ in the following way,

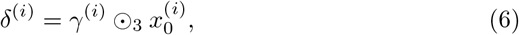

where *γ*^(*i*)^ ∈ ℝ^*c*^ defines a single factor for each channel in the image and the circled dot operator represent channel-wise multiplication, so that within each channel, the scaling factor is constant across the spatial dimensions of the image. In practice, we directly optimize for *γ*^(*i*)^, where 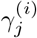 can be interpreted as the relative change to the mean intensity of the *j*-th channel. However, given our classifier does have fine spatial resolution, we can search for targeted therapies such as perturbing only a specific cell type or restricting the perturbation to specific tissue locations by changing equation (6) to match these different types of perturbation.

Taken together, the optimization procedure produces an altered image predicted to contain T cells from an original image which lacks T cells, by minimally perturbing the original image in the direction of the nearest training patch containing T cells until the classifier predicts the perturbed image to contain T cells (Figure 1C).

### Implementation of optimization procedure

We solve for the optimal perturbation *δ*^(*i*)^ for each individual patch *I*^(*i*)^ from the training cohort that 1) contain tumor cells and 2) does not contain CD8+ T cells (Figure 1C). Since our strategy may find different perturbations for different tumor patches, we reduce the set of patch-wise perturbations {*δ*^(*i*)^}_*i*_ to a whole-tumor perturbation by first taking the median across all patches for each patient, then across all patients. We evaluate the performance of a whole-tumor perturbation by applying the perturbation computationally to patches from the test cohort, before passing the perturbed patches through our trained classifier.

During optimization, the weight *c* of the loss term *L*_pred_ is updated for *n* iterations, starting at *c*_init_. If we identify a valid counterfactual (predicted to contain T cells) for the present value of *c*, we will then decrease *c* in the subsequent optimization cycle to increase the weight of the additional loss terms to help regularize our solution. If, however, we do not identify a counterfactual, *c* is increased to put more emphasis on increasing the predicted probability of the counterfactual. The parameter *s*_max_ sets the maximum number of optimization steps for each value of *c*. The parameter *l*_init_ sets the initial step size for each optimization step. Our optimization code was implemented in Python and was adapted from the Python library Alibi [57], with substantial modifications including PyTorch compatibility and improved speed.

For the purpose of speed, *L*_proto_ is defined by first building a k-d tree of training instances classified as having T cells and setting the *k*-nearest item in the tree (in terms of euclidean distance to 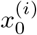 as proto. We use *k* = 1 for all counterfactual optimization. For all other parameters, we list their values in Supplemental Table 11.

### Non-spatial models

In addition to the U-Net model, we also trained a single-layer neural network on the average intensity values from each molecular channel to obtain a logistic regression classifier, predicting the probability of CD8+ T cell presence in the image patch. This model represents a linear model where only the average intensity of each molecule is used for prediction instead of their spatial distribution within a patch. Furthermore, we trained a Multilayer Perceptron (MLP) which also uses averaged intensity as input features for prediction but is capable of learning nonlinear interactions between features. The MLP model consists of two hidden layers (30 and 10 nodes) with ReLU activation.

### Primary cell isolation and cell culture

Cryopreserved human peripheral blood mononuclear cells (PBMCs; Charles River Laboratories or STEMCELL Technologies 70025.1) were thawed in RPMI media supplemented with 100 U ml^−1^ penicillin and 100 U ml^−1^ streptomycin (Thermo), and 10% fetal bovine serum (FBS; Thermo). The human melanoma cell line A375 (American Type Culture Collection, CRL-1619) was cultured in DMEM media supplemented with 100 U ml^−1^ penicillin, 100 U ml^−1^ streptomycin, 1 mM HEPES, sodium and 10% fetal bovine serum (FBS). The human colon cancer cell line HCT116 (American Type Culture Collection; kindly provided by Ekihiro Seki lab in Cedars-Sinai Medical Center) was cultured in DMEM media (10569010; Thermo) supplemented with 100 U ml^−1^ penicillin and 100 U ml^−1^ streptomycin, and 10% FBS, in an incubator at 37^°^C with 5% CO_2_.

### T-cell migration assay

We used a transwell assay to examine the impact of predicted perturbations on CD8+ T cell migration. Prior to the start of the experiment, PBMCs were thawed and rested for 12 hours, after which they were pre-treated with combinations of chemokines and antibodies for an additional 12 hours or maintained in RPMI. Tumor cells (either A375 or HCT116 cells) were thawed and grown to 80-90% confluence in the bottom chamber of 96-well plates. Cancer cells (and PBMCs for CRC) were incubated overnight in a 37^°^C incubator, with supplementation of signaling proteins and antibodies, according to Morpheus’ predictions. A full list of signaling proteins and antibodies used to implement perturbations is in Supplemental Table 12. After incubation, PBMCs were seeded into an HTS 96-well permeable support (CLS3387, Corning), which were then placed inside 96 well plates containing cancer cells. Cells were allowed to migrate for 4 hours according to published transwell migration protocols for T cells [38].

### Flow cytometric counting of CD8+ T-cells

To count CD8+ T cells, supernatants from the bottom well of the transwell assay were collected and centrifuged at 500 g for 5 minutes. The supernatant was discarded and the cell pellet was used for subsequent staining for CD8+ T cells using a mouse anti-human CD8 monoclonal antibody (CD8 Monoclonal Antibody 3B5) labeled with either Qdot 800 or FITC. PBMCs were resuspended in HBSS buffer with 10mM HEPES and 0.5% BSA. Unstained PBMCs were used as a control to determine gating strategy (Supplemental Figure 4). Staining was performed according to the manufacturer’s instructions. Flow cytometry was performed using either MACSQuant Analyzer 10/VYB (Miltenyi Biotec) or Cytoflex S Flow Cytometer (Beckman Coulter).

### Statistical analysis

We assessed the likelihood of observing a specific number of patients with a particular phenotype in a given cluster (Figure 3B, Figure 4C) using the hypergeometric test. This statistical test calculates the probability of *k* successes in *n* draws from a population of size *N* containing *m* successes, where draws are made without replacement. The formula used is,

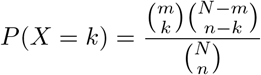

For the melanoma data set, this test was applied to determine the probability of observing the distribution of patients with stage III and stage IV melanoma among the two clusters. For the CRC data set, this test was applied to determine the probability of observing the distribution of patients with NAFLD among the two clusters.

We used the Wilcoxon rank-sum test to compare molecule levels and cell type abundance between two patient clusters using tumor tissue samples (Figure 3C, Figure 4D). This non-parametric test evaluates whether there is a significant difference in the distributions of two independent samples. The Wilcoxon rank-sum test statistic *W* is calculated as:

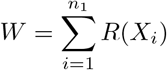

where *R*(*X*_*i*_) is the rank of the *i*-th observation from the first sample in the combined sample of size *n*_1_ + *n*_2_. To account for multiple comparisons, p-values obtained from the Wilcoxon rank-sum test were adjusted using the Šidák correction. The Šidák-adjusted p-value *p*_adj_ is given by:

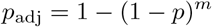

where *p* is the original p-value, and *m* is the number of tests performed. This method adjusts the p-values by calculating the cumulative probability of avoiding type I errors across all tests, providing a rigorous control of the Family-Wise Error Rate.

We used the Welsh’s t-test to assess whether there is a statistical significant difference in T cell infiltration between responder and non-responders (Supplemental Figure 3). Welsh’s t-test is used to compare the means of two independent samples without assuming equal variance. The T statistic is defined as follows,

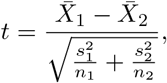

where 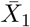 and 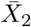 are the sample means, *s*^2^ and *s*^2^ are the sample variances, and *n*_1_ and *n*_2_ are the sample sizes.

We used paired t-tests to assess the significance of the perturbation effect across multiple replicates in our transwell assay. The paired t-test is used to compare the means of two related samples, typically before and after a treatment or intervention. The t statistic for paired samples is defined as follows:

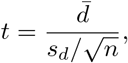

where 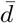 is the mean difference between paired observations, *s*_*d*_ is the standard deviation of the differences, and *n* is the number of pairs.

## Code Availability

Morpheus is available as an open-source python package at https://pypi.org/project/morpheus-spatial/. Code for model training, perturbation optimization and analysis are publicly available at https://github.com/neonine2/morpheus-spatial, which also includes a tutorial Jupyter notebook for using Morpheus. Our optimization code was implemented in Python and was adapted from the open source Python library Alibi [57], with substantia modifications.

## Data Availability

All data sets used in this study are published and publicly available. The melanoma (10.5281/zenodo.5994135) and breast tumor (10.5281/zenodo.5850951) data sets are both place in the Zenodo data repository. The CRC liver metastases is available upon request from the lead contact author (https://doi.org/10.1016/j.cmet.2023.04.013). All pre-processed data and model outputs are available via the identifier https://doi.org/10.22002/pr14s-wgk05. Jupyter notebook is available on the Github repo for reproducing primary analysis and main figures using the deposited data.

## Acknowledgements

We would like to thank Inna Strazhnik for her support with figure illustrations. We would like to thank James Linton and Michael Elowitz for support with flow cytometry workflow. We also appreciate the Ekihiro Seki lab in Cedars-Sinai Medical Center for kindly providing the HCT116 cell line. We would like to thank Akil Merchant, Aviv Regev, Long Cai, Barbara Wold, Michal Polonsky, Frederick Eberhardt and all members of the Thomson lab for insightful discussion that greatly improved this work. We gratefully acknowledge the support of the National Institutes of Health’s Information Technology for Cancer Research (ITCR) program and the Merkin Institute for Translational Research.

## Supplemental Note 1 Assessing effects of data preprocessing steps

### Effect of T cell masking on classifier

By masking T cells prior to model training, we may have inadvertently introduced a new signal that is predictive of T cells, specifically that less cells in an image increases the probability of T cells being present. To assess the possibility of such an effect, we study the impact of random cell masking on the predicted probability of T-cell presence. For each patch, we generate three randomized versions, where we randomly select one of the cells to mask for each version. We then compute the difference in predicted probabilities and predicted label between the randomized patches and the original patch (Supplemental Table 1). Across all three data sets, we do not see a statistically significant difference in the predicted labels when comparing randomly-masked patches to original patches. We do see a notable increase in the predicted probability value for CRC in the randomized images, although the mean change is very small at 1.38×10^−4^. These results suggest our T-cell masking strategy did not introduce an artificial signal whereby simply removing cells at random will increase the chance that T cells are predicted to be present.

**Supplemental Table 1:**
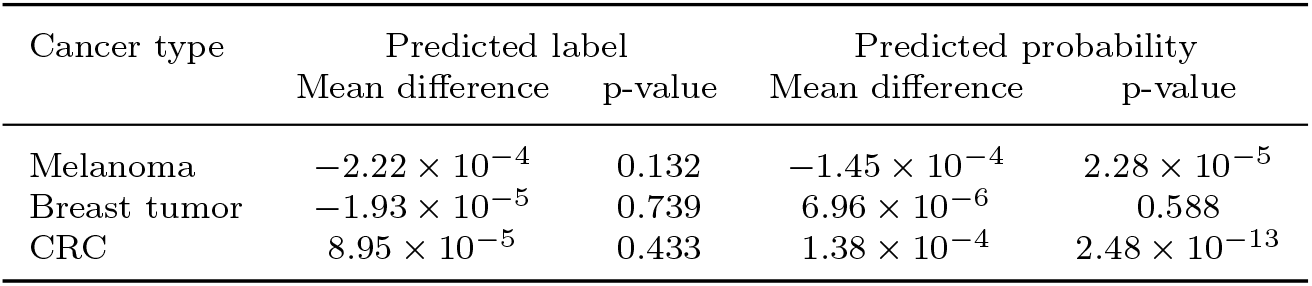
Difference in predicted probability and predicted positive labels between randomly masked patches and original patches, p-value obtained from a one-sample t-test.

| Cancer type | Predicted label |  | Predicted probability |  |
| --- | --- | --- | --- | --- |
|  | Mean difference | p-value | Mean difference | p-value |
| Melanoma | $-2.22 \times 10^{-4}$ | 0.132 | $-1.45 \times 10^{-4}$ | $2.28 \times 10^{-5}$ |
| Breast tumor | $-1.93 \times 10^{-5}$ | 0.739 | $6.96 \times 10^{-6}$ | 0.588 |
| CRC | $8.95 \times 10^{-5}$ | 0.433 | $1.38 \times 10^{-4}$ | $2.48 \times 10^{-13}$ |

### Effect of downsampling on cell count

We perform a downsampling step after cell pixelation in order to reduce the dimensionality of the input data to our classifier. Since downsampling has the potential of merging two cell pixels together, leading to loss of information, we compute the amount of cells that were merged due to downsampling and show that the loss of information is minimal as only 1 − 2% of cells were merged.

**Supplemental Table 2:** Change in cell count/pixel due to downsampling.

| Cancer type | initial cell count | cells merged | proportion merged |
| --- | --- | --- | --- |
| Melanoma | 791413 | 11171 | 1.4% |
| CRC | 926975 | 13876 | 1.5% |
| Breast tumor | 1002649 | 24840 | 2.1% |

## Supplemental Note 2 Assessing different choices of IMC patch size

In order to obtain enough TME samples to train our classifier models, we took advantage of the inherent heterogeneity in the TME and divided each tissue image into 48 µm×48 µm patches and treated each patch as an independent sample during training. Here, we perform spectral analysis to study the relationship between spatial patterning of proteins at various length scales and CD8+ T-cell infiltration. Specifically, we compute the power spectral density (PSD) of each breast tumor image. The PSD shows the relative importance of patterning at various length scales (“wavelengths”) in the expression map. We then compute the Pearson correlation between patterning of a protein at a given length scale and T-cell infiltration (Extended Data Figure 1). Extended Data Figure 1 shows there is substantial information pertaining to T-cell infiltration at our selected patch size. For certain proteins such as HLA-DR and FSP1, we see substantially more information is present at longer length scales of around 200 µm. This result suggests that for sufficient amount of IMC data, the performance of our classifier model may be improved by increasing the size of image patches.

## Supplemental Note 3 Counterfactual optimization of pseudobulk measurement

We performed additional analysis to show that single cell spatial resolution produces entirely different strategies compared to those generated with pseudobulk measurements (Extended Data Figure 2). We first sum up the expression vector across all cells in each IMC image, producing a vector of length N per image where N is the number of molecular channels. We then generated a binary label for each image by thresholding the total number of T-cells in the image, where the threshold is set to the 95th percentile of the T-cell counts from the training images. We then trained a 2-layer MLP to predict this binary label using the bulk protein measurements, followed by counterfactual optimization using the trained model. For both melanoma and CRC, the bulk level strategy is very different compared to the patch level which Morpheus operates on. For melanoma (Extended Data Figure 2A), the bulk-level strategy mainly involves increasing CXCL10 and inhibiting CCL8, with no changes to CCL18 or CCL22 which are two important targets discovered by Morpheus. For CRC (Extended Data Figure 2B), the bulk-level strategy consists of increasing IL10 while inhibiting Galectin9 and PDL1, which is again very different from the Morpheus strategy. Also, it is not clear that increasing IL10 is an effective strategy as it is known to be an anti-inflammatory cytokine.

**Supplemental Figure 1 Morpheus output for primary CRC and liver metastases**

**Supplemental Figure 1:**
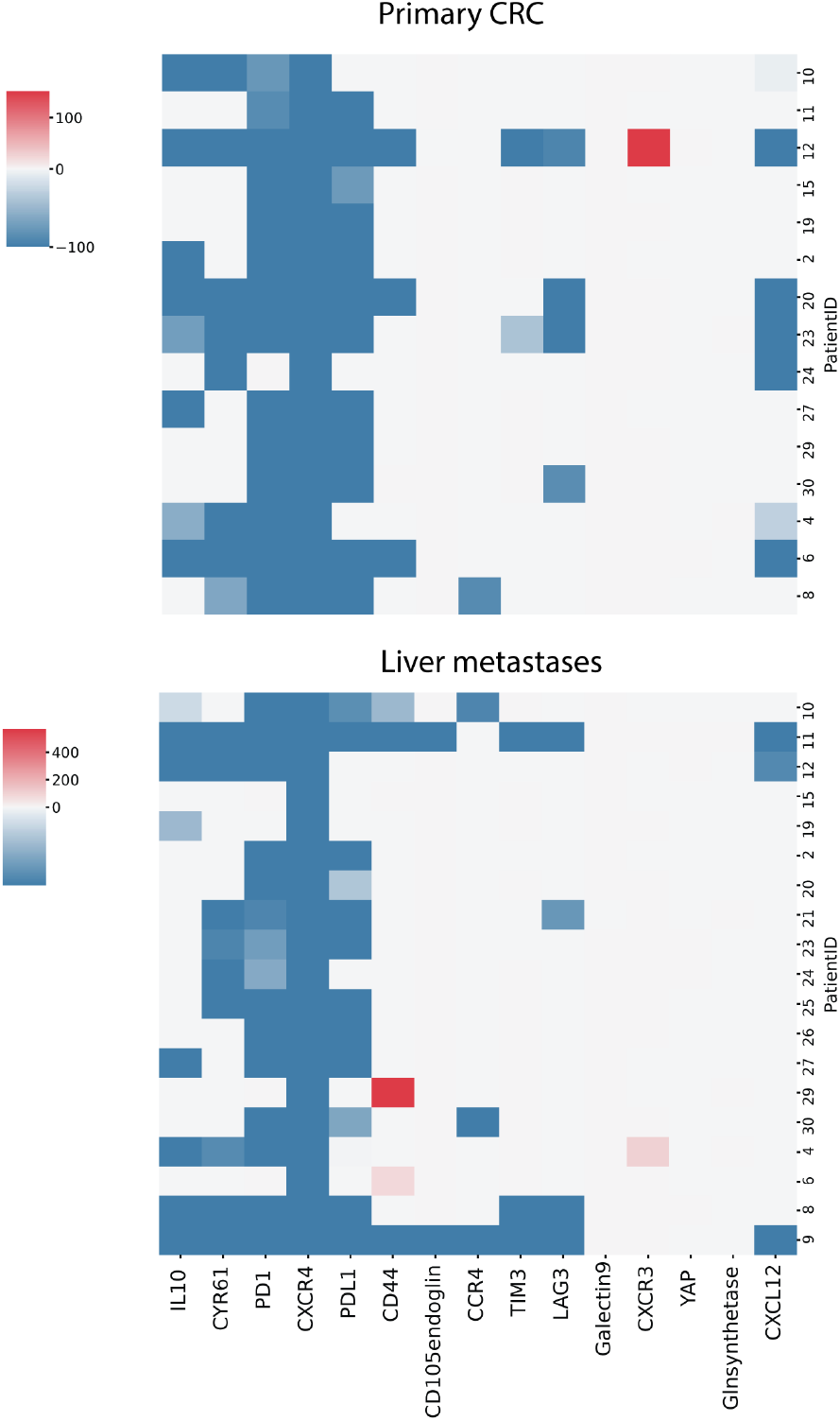
Optimized perturbations for tissue sections from primary colorectal tumor and liver metastases aggregated to the patient (row) level (train cohort).

**Supplemental Figure 2 Receiver Operating Characteristic Comparison**

**Supplemental Figure 2:**
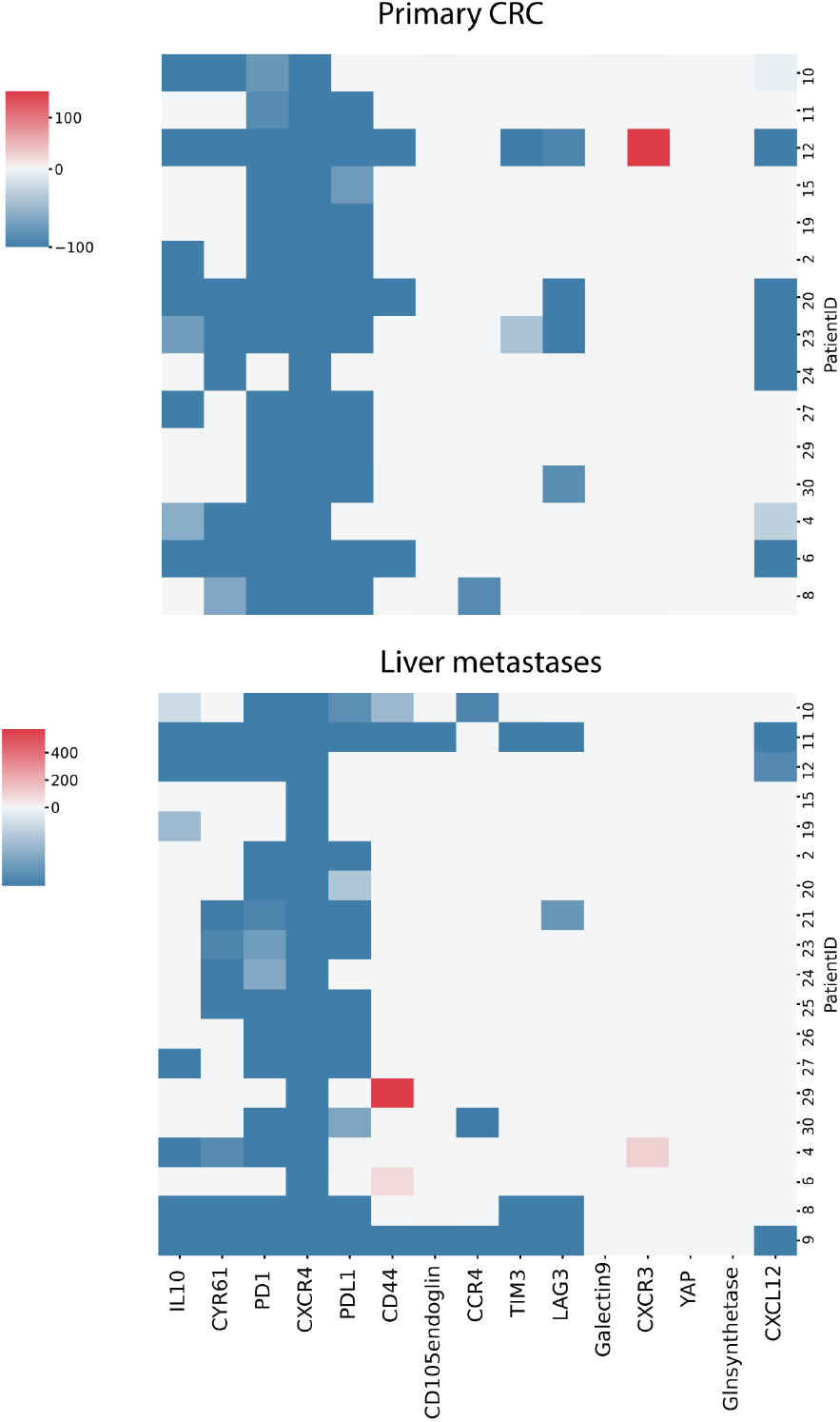
ROC curves for U-NET models trained on the melanoma, CRC and breast tumor data set. Area indicates the Area Under the ROC Curve (AUROC).

**Supplemental Table 3 Cell type distribution across data sets**

**Supplemental Table 3:** Top eight cell types by abundance across all three data sets.

| Metastatic melanoma |  | CRC |  | Breast tumor |  |
| --- | --- | --- | --- | --- | --- |
| Cell type | Count | Cell type | Count | Cell type | Count |
| Tumor | 548958 | Tumor | 287006 | Tumor | 601652 |
| Macrophage | 64408 | Liver | 249571 | Fibroblasts | 122641 |
| Cytotoxic T cell | 36722 | Stromal cell | 192217 | Myofibroblasts | 91066 |
| HLA-DR+ cells | 29557 | MDSC | 41967 | Endothelial | 48508 |
| Stromal cell | 24702 | CD4+ T cell | 30749 | Macrophages | 40585 |
| Endothelial cell | 21297 | Macrophage | 30531 | Cytotoxic T cell | 34588 |
| CD38+ cell | 11123 | Dendritic cell | 24347 | B cells | 28341 |
| Neutrophil | 5018 | Cytotoxic T cell | 23440 | CD4+ T cell | 15647 |

**Supplemental Figure 3 Relationship between T-cell infiltration level and patient response**

**Supplemental Figure 3:**
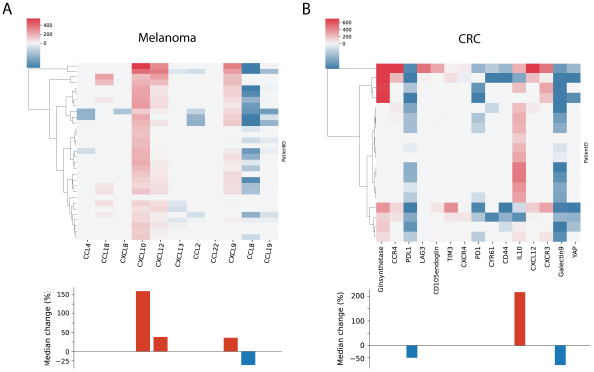
T-cell infiltration level in IMC images of patients with melanoma taken pre-treatment, grouped by whether the patient responded to later drug treatments. Response data obtained at 3 month, where responders consist of complete and partial response, and nonresponders consist of progressive disease and stable disease. Treatments include checkpoint inhibitors and chemotherapy drugs. Welch’s t-test performed to assess difference between nonresponder and responder, T-statistic = 2.95, p = 0.006

**Supplemental Table 4 Evaluation metric for all trained classifiers**

**Supplemental Table 4:**
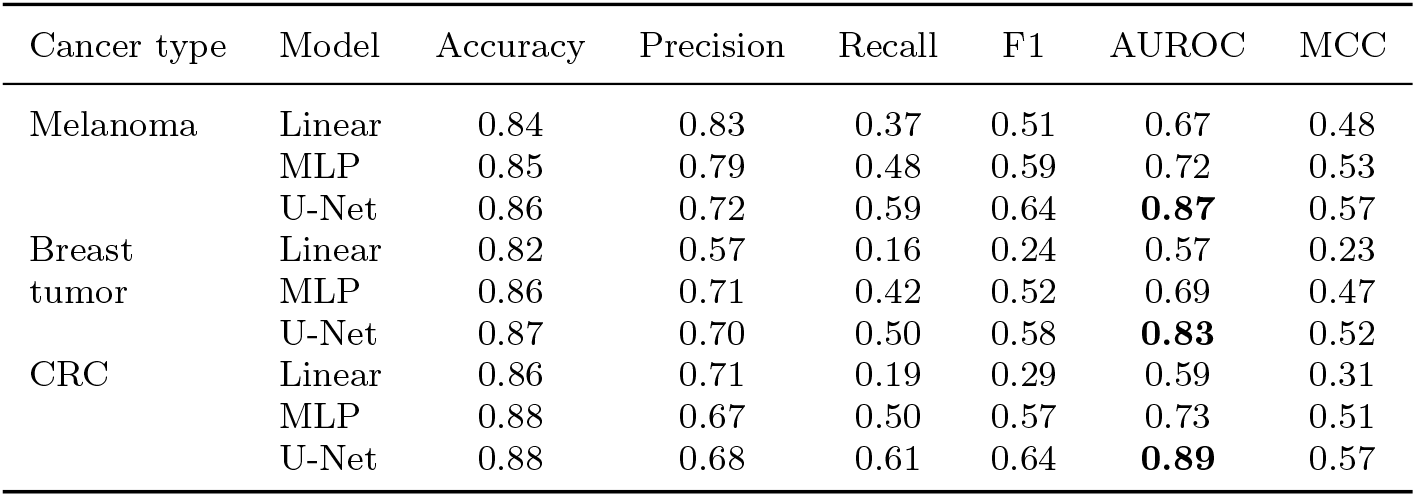
Performance of different classifier models trained on melanoma, CRC, and breast tumor IMC images to predict the presence of T cells (*p* = 0.5)

| Cancer type | Model | Accuracy | Precision | Recall | F1 | AUROC | MCC |
| --- | --- | --- | --- | --- | --- | --- | --- |
| Melanoma | Linear | 0.84 | 0.83 | 0.37 | 0.51 | 0.67 | 0.48 |
|  | MLP | 0.85 | 0.79 | 0.48 | 0.59 | 0.72 | 0.53 |
|  | U-Net | 0.86 | 0.72 | 0.59 | 0.64 | **0.87** | 0.57 |
| Breast tumor | Linear | 0.82 | 0.57 | 0.16 | 0.24 | 0.57 | 0.23 |
|  | MLP | 0.86 | 0.71 | 0.42 | 0.52 | 0.69 | 0.47 |
|  | U-Net | 0.87 | 0.70 | 0.50 | 0.58 | **0.83** | 0.52 |
| CRC | Linear | 0.86 | 0.71 | 0.19 | 0.29 | 0.59 | 0.31 |
|  | MLP | 0.88 | 0.67 | 0.50 | 0.57 | 0.73 | 0.51 |
|  | U-Net | 0.88 | 0.68 | 0.61 | 0.64 | **0.89** | 0.57 |

**Supplemental Table 5 Performance of different CNN and ViT model architectures**

**Supplemental Table 5:**
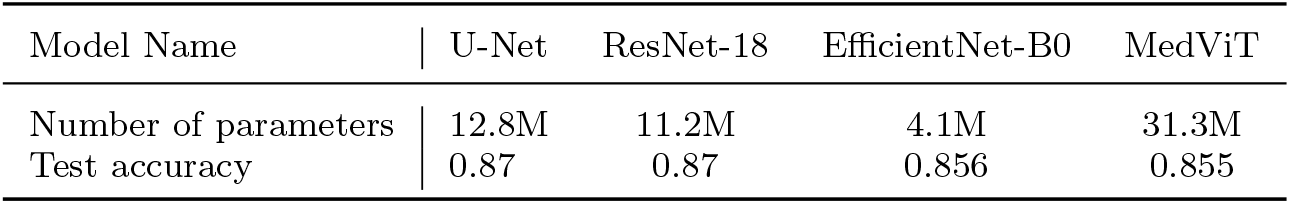
Performance of different CNN and ViT models trained on IMC image patches of melanoma.

| Model Name | U-Net | ResNet-18 | EfficientNet-B0 | MedViT |
| --- | --- | --- | --- | --- |
| Number of parameters | 12.8M | 11.2M | 4.1M | 31.3M |
| Test accuracy | 0.87 | 0.87 | 0.856 | 0.855 |

**Supplemental Figure 4 Gating Strategies for analyzing T-cell migration**

**Supplemental Figure 4:**
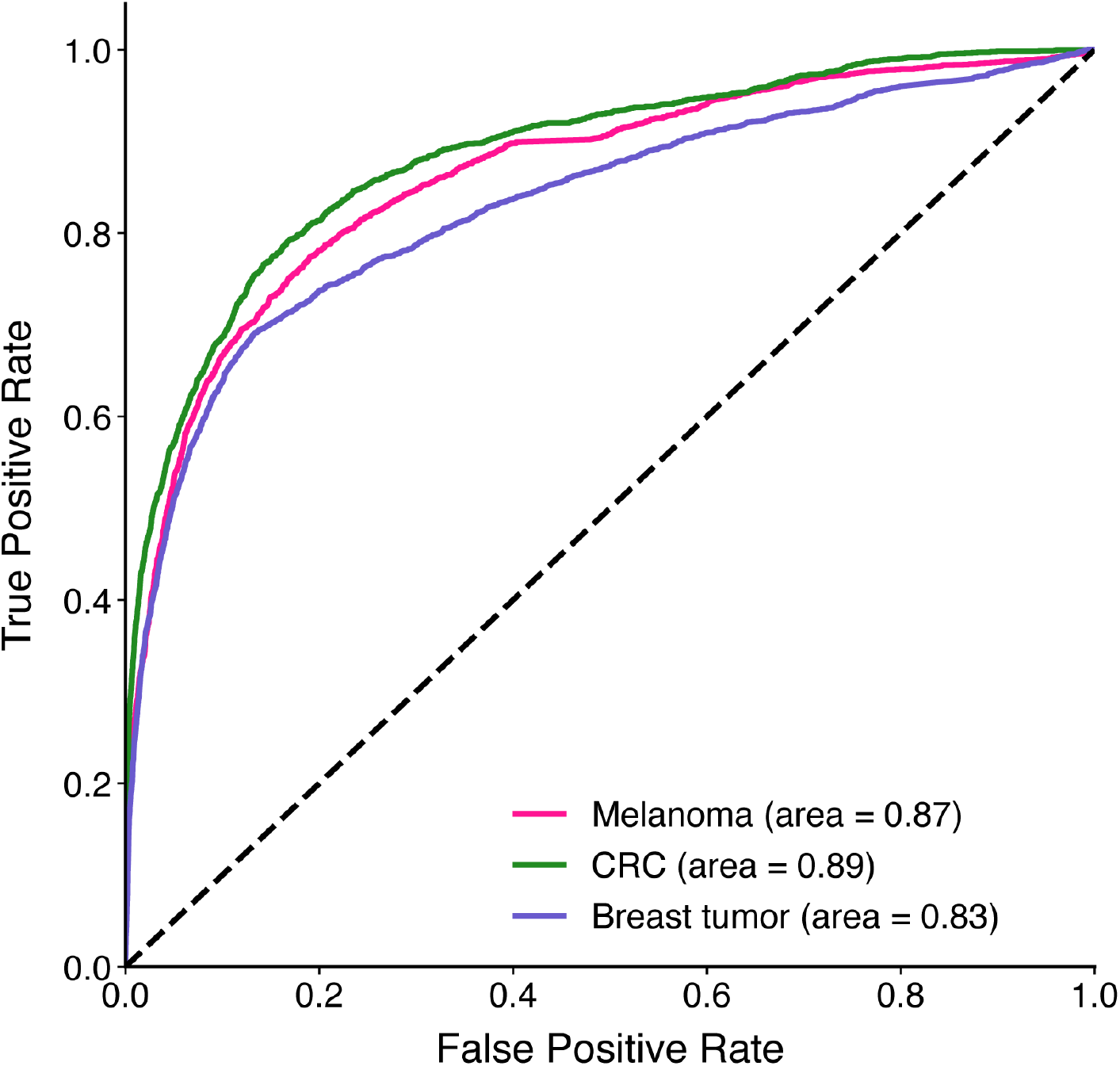
Gating strategy for counting CD8+ T cells following transwell migration. Live cells were gated to exclude doublets and debris using SSC and FSC. Thresholds for FITC positivity were determined using unstained controls.

**Supplemental Table 6 and 7 Source Data for Figure 5**

**Supplemental Table 6:**
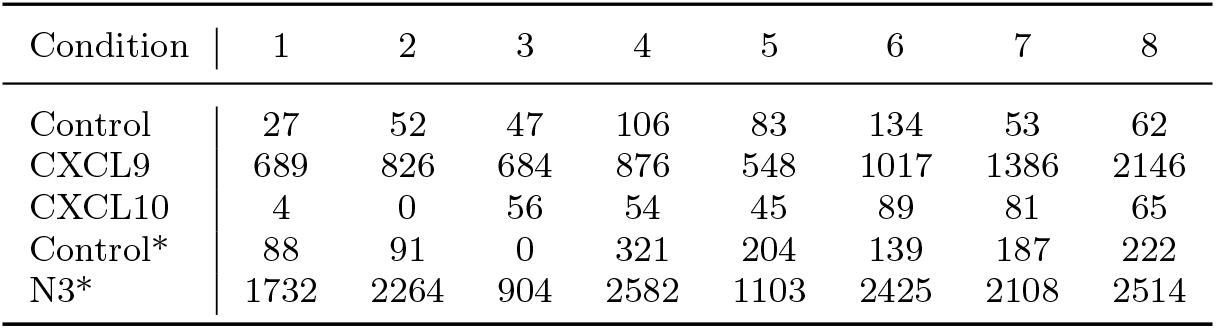
Raw CD8+ T-cell count for different experimental conditions of the transwell assay using A375 human melanoma cell line. N3: CXCL10 + *α*CCL22 + *α*CCL18. Star indicates N3 and the second set of control are part of the same experiment.

| Condition | 1 | 2 | 3 | 4 | 5 | 6 | 7 | 8 |
| --- | --- | --- | --- | --- | --- | --- | --- | --- |
| Control | 27 | 52 | 47 | 106 | 83 | 134 | 53 | 62 |
| CXCL9 | 689 | 826 | 684 | 876 | 548 | 1017 | 1386 | 2146 |
| CXCL10 | 4 | 0 | 56 | 54 | 45 | 89 | 81 | 65 |
| Control* | 88 | 91 | 0 | 321 | 204 | 139 | 187 | 222 |
| N3* | 1732 | 2264 | 904 | 2582 | 1103 | 2425 | 2108 | 2514 |

**Supplemental Table 7:** Raw CD8+ T-cell count for different experimental conditions of the transwell assay using HC116 human colon carcinoma cell line. N3: *α*PD-1 + *α*PD-L1 + *α*CXCR4, N4: *α*PD-1 + *α*PD-L1 + *α*CXCR4 + *α*CYR61.

| Condition \ Trial | 1 | 2 | 3 | 4 | 5 | 6 | 7 | 8 | 9 | 10 | 11 | 12 |
| --- | --- | --- | --- | --- | --- | --- | --- | --- | --- | --- | --- | --- |
| Control | 14783 | 22741 | 16778 | 7837 | 13766 | 10485 | 7323 | 5477 | 6880 | 1703 | 869 | 236 |
| N3 | 18632 | 24836 | 17161 | 21362 | 11674 | 22670 | 7078 | 5738 | 6376 | 3494 | 1004 | 610 |
| N4 | 26956 | 31953 | 23624 | 24540 | 21443 | 19104 | 9750 | 7734 | 7202 | 6027 | 1572 | 762 |
| αPD-1 | 20464 | 22353 | 23738 | 22664 | 8884 | 19083 | 4602 | 7338 | 5224 | | | |
| αPD-L1 | 23392 | 16974 | 22157 | 20932 | 16915 | 17898 | 9233 | 6226 | 6424 | | | |

**Supplemental Table 8 Molecular panels for datasets used**

**Supplemental Table 8:** Protein and RNA panels imaged for each of the IMC data sets, with RNA targets bolded.

| Metastatic melanoma |  | CRC with liver metastases |  | Breast tumor |  |
| --- | --- | --- | --- | --- | --- |
| Vimentin | **DapB** | CD45 | Glnsynthetase | Histone H3 | SMA |
| CD163 | **CCL4** | CD163 | NKG2D | CK5 | CD38 |
| B2M | **CCL18** | CCR4 | PD-L1 | HLA-DR | CK8-18 |
| CD134 | **CXCL8** | FAP | CD11c | CD15 | FSP1 |
| CD68 | **CXCL10** | LAG3 | HepPar1 | CD163 | ICOS |
| GLUT1 | **CXCL12** | FOXP3 | αSMA | OX40 | CD68 |
| CD3 | **CXCL13** | CD4 | CD105 | HER2 (3B5) | CD3 |
| LAG3 | **CCL2** | CD68 | VISTA | Podoplanin | CD11c |
| PD-1 | **CCL22** | CD20 | CD8α | PD-1 | GITR |
| HistoneH3 | **CXCL9** | TIM3 | CXCR4 | CD16 | c-Caspase3 |
| CCR2 | **CCL19** | PD-1 | iNOS | CD45RA | B2M |
| PD-L1 | **CCL8** | CD31 | CYR61 | CD45RO | FOXP3 |
| CD8 | SMA | CDX2 | CAIX | CD20 | ER |
| SOX10 | CD31 | CD3 | CD44 | CD8 | CD57 |
| Mart1 | pRB | CD15 | CD11b | Ki-67 | PDGFRβ |
| cleavedPARP | MPO | HLA-DR | IL10 | Caveolin-1 | CD4 |
| CD15 | CK5 | CXCL12 | HLA-ABC | CD31-vWF | CXCL12 |
| CD38 | HLA-DR | GranzymeB | Ki67 | HLA-ABC | panCK |
| S100 | Cadherin11 | HistoneH3 | CXCR3 | HER2 (D8F12) |  |
| FAP |  | Galectin9 | YAP |  |  |
|  |  | CD14 | CK19 |  |  |

**Supplemental Table 9 Data splits**

**Supplemental Table 9:** Data split for Melanoma, CRC cohort, and breast tumor IMC data set.

| Data set | Group | Patient count | Patch count | Proportion of patches<br>with CD8+ T cells |
| --- | --- | --- | --- | --- |
| Metastatic<br>melanoma | Training | 102 | 23741 | 29.6% |
|  | Validation | 28 | 6045 | 30.3% |
|  | Testing | 29 | 5950 | 30.4% |
| CRC with<br>liver metastases | Training | 19 | 44449 | 15.9% |
|  | Validation | 4 | 6957 | 14.4% |
|  | Testing | 7 | 14907 | 15.9% |
| Breast cancer | Training | 485 | 41104 | 23.7% |
|  | Validation | 113 | 9015 | 23.4% |
|  | Testing | 151 | 12987 | 23.8% |

**Supplemental Table 10 Cell phenotyping markers**

**Supplemental Table 10:** proteins for identification of T cells and tumor cells.

| Cell type | Proteins |
| --- | --- |
| CD8+ T cell | CD8, CD3 |
| Tumor cell (Melanoma) | SOX9, SOX10, MITF, Mart1, S100A1, p75 |
| Tumor cell (CRC and liver metastasis) | YAP1, CDX-2 |
| Tumor cell (breast tumor) | ER, HER2, CD15, CD57 |

**Supplemental Table 11 Parameters for counterfactual optimization**

**Supplemental Table 11:** Parameter values used for counterfactual optimization.

| Parameters | Melanoma | CRC |
| --- | --- | --- |
| $\beta$ | 2 | 50 |
| $\theta$ | 60 | 80 |
| $\kappa$ | -0.7 | -0.34 |
| $c_{\text{init}}$ | 25000 | 1000 |
| $n$ | 5 | 5 |
| $s_{\text{max}}$ | 1000 | 1000 |
| $l_{\text{init}}$ | 0.1 | 0.1 |

**Supplemental Table 12 Reagents for transwell assay**

**Supplemental Table 12:** Signaling proteins and antibodies reagents used for perturbation experiments.

| Catalog Number | Type of protein | Name | Melanoma/CRC perturbation |
| --- | --- | --- | --- |
| MAB10861,<br>R&D Systems | Antibody | Human anti-PD-1<br>Antibody | CRC |
| PA1-16579,<br>Thermo | Antibody | Human anti-CYR61<br>Polyclonal Antibody | CRC |
| MIH1, Thermo | Antibody | Human anti-PD-L1<br>Monoclonal Antibody | CRC |
| MAB170,<br>R&D Systems | Antibody | Human anti-CXCR4<br>Antibody | CRC |
| AB9849,<br>Abcam | Antibody | Human anti-CCL18<br>antibody | melanoma |
| MAB336,<br>R&D Systems | Antibody | Human anti-CCL22<br>antibody | melanoma |
| AB283902,<br>Abcam | Chemokine | Recombinant human<br>CXCL9 | melanoma |
| AB280332,<br>Abcam | Chemokine | Recombinant human<br>CXCL10 | melanoma |

